# Leaflet specific phospholipid imaging using genetically encoded proximity sensors

**DOI:** 10.1101/2024.05.01.592120

**Authors:** William M. Moore, Roberto J. Brea, Caroline H. Knittel, Ellen Wrightsman, Brandon Hui, Jinchao Lou, Christelle F. Ancajas, Michael D. Best, Christopher J. Obara, Neal K. Devaraj, Itay Budin

## Abstract

The lipid composition of cells varies widely across organelles and between individual membrane leaflets. Transport proteins are thought to generate this heterogeneity, but measuring their functions *in vivo* has been hampered by limited tools for imaging lipids at relevant spatial resolutions. Here we present fluorogen-activating coincidence encounter sensing (FACES), a chemogenetic tool capable of quantitatively imaging subcellular lipid pools and reporting their transbilayer orientation in living cells. FACES combines bioorthogonal chemistry with genetically encoded fluorogen-activating proteins (FAPs) for reversible proximity sensing of conjugated molecules. We first apply this approach to identify roles for lipid transfer proteins that traffic phosphatidylcholine pools between the ER and mitochondria. We then show that transmembrane domain-containing FAPs can reveal the membrane asymmetry of multiple lipid classes in the trans-Golgi network and be used to investigate the mechanisms that generate it. Lastly, we demonstrate FACES can be applied to measure other molecule classes, like sugars.

## Introduction

Lipid composition determines the biophysical properties of cell membranes and varies across every scale of biological organization^1–3^. In cells, membrane heterogeneity between organelles is generated by a complex network of lipid synthesis and transport pathways that can become concentrated at membrane contact sites; hot zones of lipid transport where opposing organelles are in close proximity, often only tens of nm apart^4,5^. While many lipid transfer proteins (LTP) acting at these sites have now been biochemically characterized^6,7^, measuring their in vivo functions remains challenging. Within organelles, it has long been recognized that each bilayer leaflet – separated by less than 4 nm – could feature a unique lipidome, which is reflected in the biophysical properties of each leaflet and the topology of protein transmembrane domains^8^. Detailed lipidomic investigation of membrane asymmetry has only been carried out in the PM of erythrocytes, whose lack of intracellular compartments allow for calculation of lipid accessibility to externally added phospholipases^3,9^. While asymmetry is an emerging concept in membrane biology, the transbilayer distribution of phospholipids in intracellular membranes and its molecular drivers are still poorly understood.

Investigating membrane chemical heterogeneity remains a challenge due to the paucity of tools for imaging lipids in live cells at spatial resolutions needed to investigate lipid transport or asymmetry. Bioorthogonal ‘click’ chemistry has emerged as one tool for fluorescence labeling of specific lipid classes – including phosphatidylcholine (PC)^10^, phosphatidylserine (PS)^11^, phosphatidylinositol (PI)^12^, phosphatidic acid^13^, and glycosphingolipids^14^ – that can be metabolically tagged with azide-bearing head group components. Azido (N_3_) lipids are commonly conjugated to fluorophores using strain-promoted azide-alkyne cycloaddition (SPAAC) reactions, unveiling their cellular distribution once unreactive dyes are washed out. While powerful, this approach indiscriminately labels all lipids in the cell, and is thus limited by the resolution constraints of light microscopy; it cannot discriminate between lipid pools in organelles at close proximity or in individual leaflets of the same membrane, for example. To overcome some of these limitations, organelle-targeted fluorophores have been applied to image subcellular lipid pools^15^, but their specificity is limited to just a few chemical environments in the cell.

We hypothesized that challenges for lipid imaging could be overcome using protein-based lipid sensors, whose localization is precisely controlled through signal sequences or fusion to transmembrane domains. Previously, natural lipid binding proteins derived from microbial toxins, such as perfringolysin O^16^ (cholesterol) or equinatoxin II^17^ (sphingomyelin, SM) have been expressed as fusions with fluorescent proteins (FP), to detect membranes enriched in the target lipids. In these modalities, lipids are only qualitatively detected by colocalization of a probe to a membrane, which limits their application to large, well-separated compartments. We asked whether fluorogenic systems could provide a more generalizable strategy. Fluorogen-activating proteins (FAPs) can be expressed as fusion constructs with other proteins or targeting sequences, similar to FPs, but are not themselves fluorescent^18^. Instead, they interact with fluorogens, small cell-permeable molecules that act as fluorophores only upon specific binding to a FAP. Depending on the FAP used, fluorogen binding can be irreversible or reversible; the former provides brighter fluorescence for conventional imaging, while the latter generates stochastic blinking that can be harnessed for super-resolution imaging^19^. Recently, FAPs have been applied towards detecting and controlling protein proximity in living cells through reversible dimerization^20^, demonstrating that this technology can be harnessed in new applications.

Here, we seek to harness the unique properties of fluorogens through Fluorogen-Activating Coincidence Encounter Sensing (FACES), a method that uses FAPs for proximity-dependent reporting of fluorogen conjugated bioorthogonal metabolites, including lipids. We apply the FACES strategy to investigate the transport of N_3_-phospholipids in cells and characterize the role of ER-mitochondrial PC transporters. We then show that transmembrane-anchored FAPs can probe leaflet-specific membrane composition, identifying features and drivers of asymmetry generated during vesicle formation at the trans-Golgi network (TGN). Lastly, we demonstrate that FACES is a broadly applicable tool compatible with non-lipid azido-labeled metabolites by measuring fluctuations in mitochondrial *N*-acetylhexosamine (HexNAz) levels.

## Results

### Development and validation of FACES

To implement FACES, we sought to couple fluorogenic dyes to label azido-labeled metabolites that incorporate in native phospholipid metabolism (Fig. 1a). The conjugated fluorogen tags would then only fluoresce if bound to a FAP (Fig. 1b). To accomplish this, we synthesized derivatives of the fluorogen malachite green (MG) with a linker connected to a dibenzocyclooctyne (DBCO) group for copper-free SPAAC reactions, including compound **1a** (Fig. 1c; Supplementary Fig. 1). MG non-covalently binds to soluble single-chain antibody scFvs FAP and the complex emits far-red fluorescence. In the FACES workflow (Fig. 1d), cells first express a FAP fused to an FP, like GFP, that is targeted to an organelle or site of interest. Cells are then incubated with azido-labeled metabolites for metabolic incorporation. Immediately before imaging, incorporated metabolites are labeled with fluorogen-DBCO, which remains non-fluorescent unless it is bound to the expressed FAP. Excess dye is removed from cells by washing with culture medium, leveraging the reversibility of FAP binding to minimize detection of unreacted fluorogens. Fluorogenic fluorescence thus acts like an AND-gate: it requires both the target molecule be modified with a MG green molecule, and the corresponding FAP to be in the same compartment or membrane. The abundance of the target molecule is measured with the ratio of fluorogen fluorescence, which is at red shifted wavelengths for MG, to that of the reporter FP (e.g. GFP). This ratiometric read out normalizes the concentration of the FAP receptor itself.

**Figure 1.**
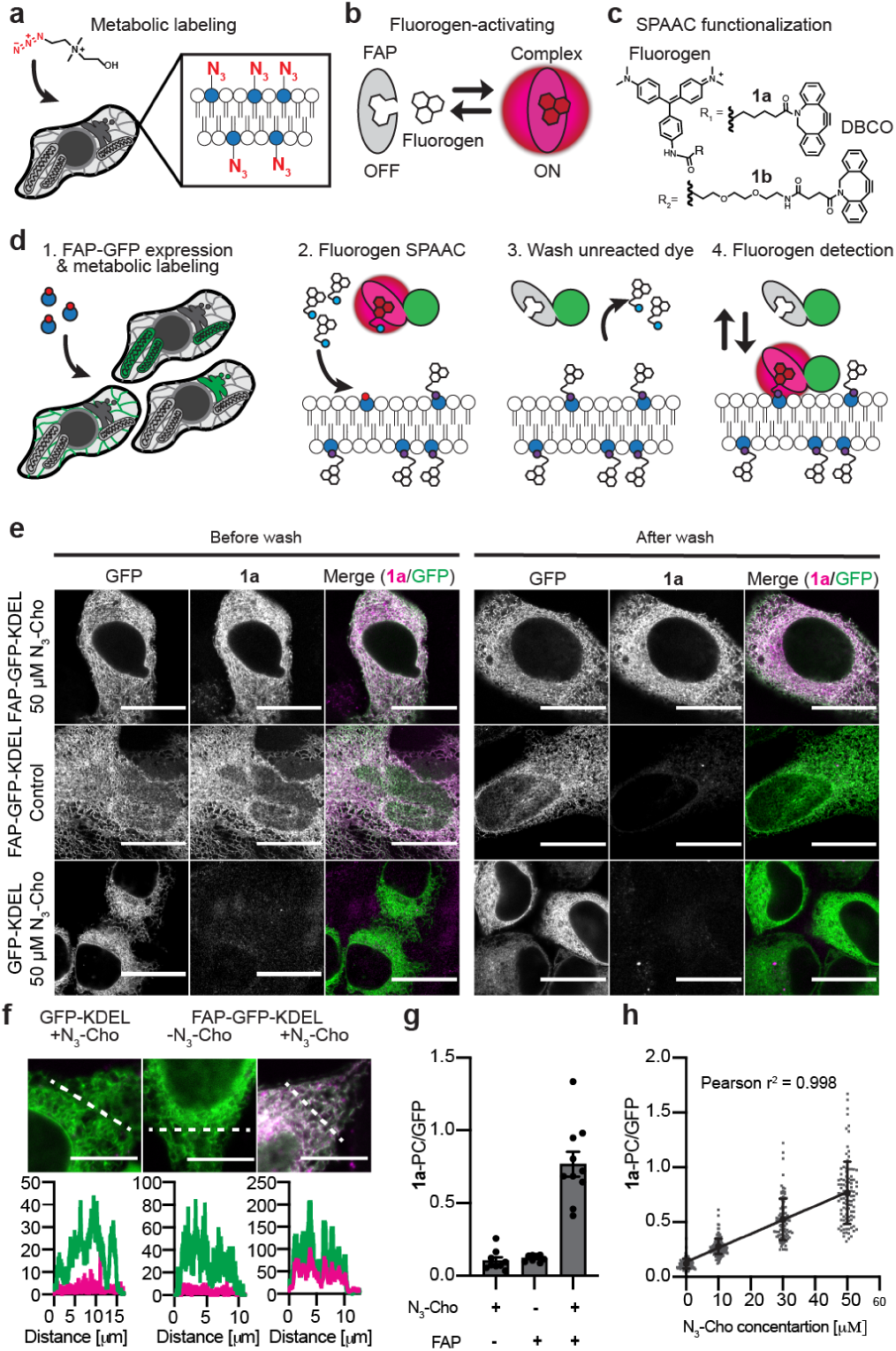
Conceptual development of FACES **a,** 1-azidoethyl-choline (N_3_-Cho) is metabolically incorporated into azido-PC lipids distributed throughout all cellular membranes. **b,** Fluorogens form reversible non-covalent fluorescent complexes with genetically encoded FAPs. **c,** Structure of the fluorogen malachite green linked to dibenzocyclooctyne (DBCO) for SPAAC labeling. **d,** Conceptual schematic detailing the FACES method. (1) Cells expressing a FAP-GFP fusion protein targeted to an organelle of interest are metabolically labeled with N_3_-Cho, or other bioorthogonal metabolite. (2) After incorporation, metabolites are conjugated via a SPAAC reaction with fluorogen-DBCO. During this time free dye can interact with the FAP to generate far-red fluorescence. (3) After SPAAC labeling, excess free dye is reversibly disassociated from the FAP complex and removed from cells by washing. (4) Subsequent imaging of the fluorogen-FAP complex is used to detect the specific biomolecule pool. Co-measurement of GFP fluorescence is used to normalize the fluorogen signal to FAP sensor density. **e,** HeLa cells transfected with either FAP-GFP-KDEL (top two rows) or GFP-KDEL (bottom row) were grown in the presence (top and bottom rows) or absence (middle row) of 50 μM N_3_-Cho and labeled with 500 nM **1a**. Cells were imaged while in the labeling solution (left) and after washing with culture medium (right). Scale = 20 μm. **f,** Merged confocal images of transfected HeLa cells post washing. White line indicates the segment used for plotting the fluorescent intensity profile of **1a** and GFP shown below. Scale = 10 μm. **g,** Quantified fluorescence intensity profiles from confocal images expressed as ratiometric average; *n = 1, N* = 10 cells. Error bars reflect SEM. **h,** Linear regression of quantified fluorescence from transfected HeLa cells fed 0, 10, 30, 50 μM N_3_-Cho and labeled with 500 nM **1a**;*n* = 1, *N* = 100 individual ER segments derived from 10 cells. Error bars reflect SD.

We tested the compatibility and fluorogenic properties of **1a** with FACES using the FAP clone dH6.2, a lower affinity scFvs variant developed for reversible equilibrium binding to MG^21^. In all experiments, we fused dH6.2 – from here on simply referred to as FAP – to GFP for ratiometric imaging. An N-terminal secretion signal peptide and C-terminal KDEL retention sequence were first used to target FAP-GFP to the endoplasmic reticulum (ER) as the major site of PC synthesis^22^, while GFP-KDEL, an identical construct but lacking the FAP domain, was used as a control. HeLa cells transfected with either FAP-GFP-KDEL or GFP-KDEL were grown for 24h in the presence or absence of 50 μM 1-azidoethyl-choline (N_3_-Cho). At concentrations up to 100 μM, N_3_-Cho was incorporated into PC at a frequency up to 0.1 mol % of unlabeled PC, but cells showed no detectable N_3_-Cho incorporation into SM (Supplementary Fig. 2). Prior to staining, cells were washed and grown in normal growth media for 30 min to allow for the metabolic consumption of any remaining N_3_-Cho. Cells were then transferred into FBS-free growth media containing 500 nM **1a** and imaged while in the staining solution (Fig. 1e). Images were collected using identical acquisition and processing settings.

Results from pre-washed cells demonstrated that **1a** was cell permeable and generated strong far-red fluorescence in cells expressing FAP-GFP-KDEL (Fig. 1e). No far-red fluorescence was detected in cells expressing GFP-KDEL, indicating that **1a** was not activated directly by conjugation to N_3_-PC. Next we tested the reversibility of the FAP-fluorogen complex by washing cells with growth medium and re-imaging. Three washes were sufficient to dissociate free **1a** from FAP-GFP-KDEL cells in the absence of N_3_-Cho, resulting in the loss of far-red fluorescence. In contrast, far-red fluorescence was retained in FAP-GFP-KDEL cells that were fed 50 μM N_3_-Cho, which implicated the SPAAC reaction product **1a**-PC as the source of FAP-dependent fluorescence (Fig. 1e). Similar experiments with a higher affinity FAP variant retained far-red fluorescence after washing in the absence of N_3_-Cho labeling (Supplementary Fig. 3), indicating that the reversible binding to the MG product is essential for substrate-specific detection. In this regard, each experiment necessitates that a no azido control be included to assess the wash out of fluorogen and measure to the intrinsic background fluorescence.

We further quantified these results in repeated experiments using confocal laser scanning microscopy (CFLSM) to assess linear fluorescent detection. Cells transfected with FAP-GFP-KDEL were fed 10, 30, or 50 μM N_3_-Cho for 24h, while FAP-GFP-KDEL (no N_3_-Cho) and GFP-KDEL (+ 50 μM N_3_-Cho) conditions were used as controls. All cells were stained with 500 nM **1a** and were washed as previously described; the **1a**-PC reaction product was confirmed by LC-MS/MS (Supplementary Fig. 4). Fluorescence intensity profiles revealed that far-red fluorescence from **1a** was tightly correlated with GFP fluorescence in FAP-GFP-KDEL cells fed 50 μM N_3_-Cho (Fig. 1f). In contrast, cells from either control condition, lacking the FAP or N_3_-Cho component, had low levels of far-red fluorescence that did not correlate with GFP localization. The average fluorescence intensity from 10 cells, using 10 individual ER segments per cell, was calculated as a ratiometric average relative to GFP (Fig. 1g). There was no intensity difference between either control condition, while signal from FAP-GFP-KDEL cells fed 50 μM N_3_-Cho was 10-fold higher. Additionally, **1a**-PC fluorescence was linearly dependent on N_3_-Cho concentration in the medium (Fig. 1h). Together, these results indicate that FACES is capable of linear fluorescent quantification of N_3_-PC, via its SPAAC reaction product **1a**-PC, in the ER of living cells.

### Imaging suborganellar PC distribution in mitochondria

To expand FACES to other organelles we used the OTC mitochondrial targeting sequence (MTS) to direct FAP-GFP to the mitochondrial matrix. When quantifying the performance of FACES in the mitochondrial matrix using CFLSM, we found that switching the C_6_ alkane linker in **1a** to a polyethylene glycol (PEG)-2 linker (compound **1b**) was beneficial for dissociating unreacted dye from this compartment, reducing background fluorogen fluorescence in the absence of the SPAAC reaction. We suspect that the denser membranes of the IMM and/or presence of the outer mitochondrial membrane as an additional retention barrier, necessitating the more polar fluorogen **1b** for optimal performance. Similar to the ER-targeted protein, MTS-FAP-GFP detection of **1b**-PC generated fluorescence that linearly correlated with N_3_-Cho concentration (Supplementary Fig.5a-c).

Using super-resolution lattice-structured illumination microscopy (lattice SIM^2^), we further tested whether the matrix soluble FAP could resolve structural features of the IMM. When imaged by lattice SIM^2^, FACES signal from **1b**-PC occurred in a banding pattern distinct from the general distribution of GFP in the mitochondrial matrix (Supplementary Fig. 5d). These experiments were repeated with cells counterstained with PKmito^23^ to label mitochondrial cristae (Fig. 2a). Fluorescence intensity profiles drawn across individual mitochondria show that **1b**-PC correlated well with PKmito, which at times formed alternating peaks with GFP (Fig. 2b). We interpreted this localization pattern as reflecting regions of high cristae density in the IMM that contrast with void areas of the matrix^24^. This is in line with previous work using a chemically targeted approach to label mitochondrial N_3_-PC^15^.

**Figure 2.**
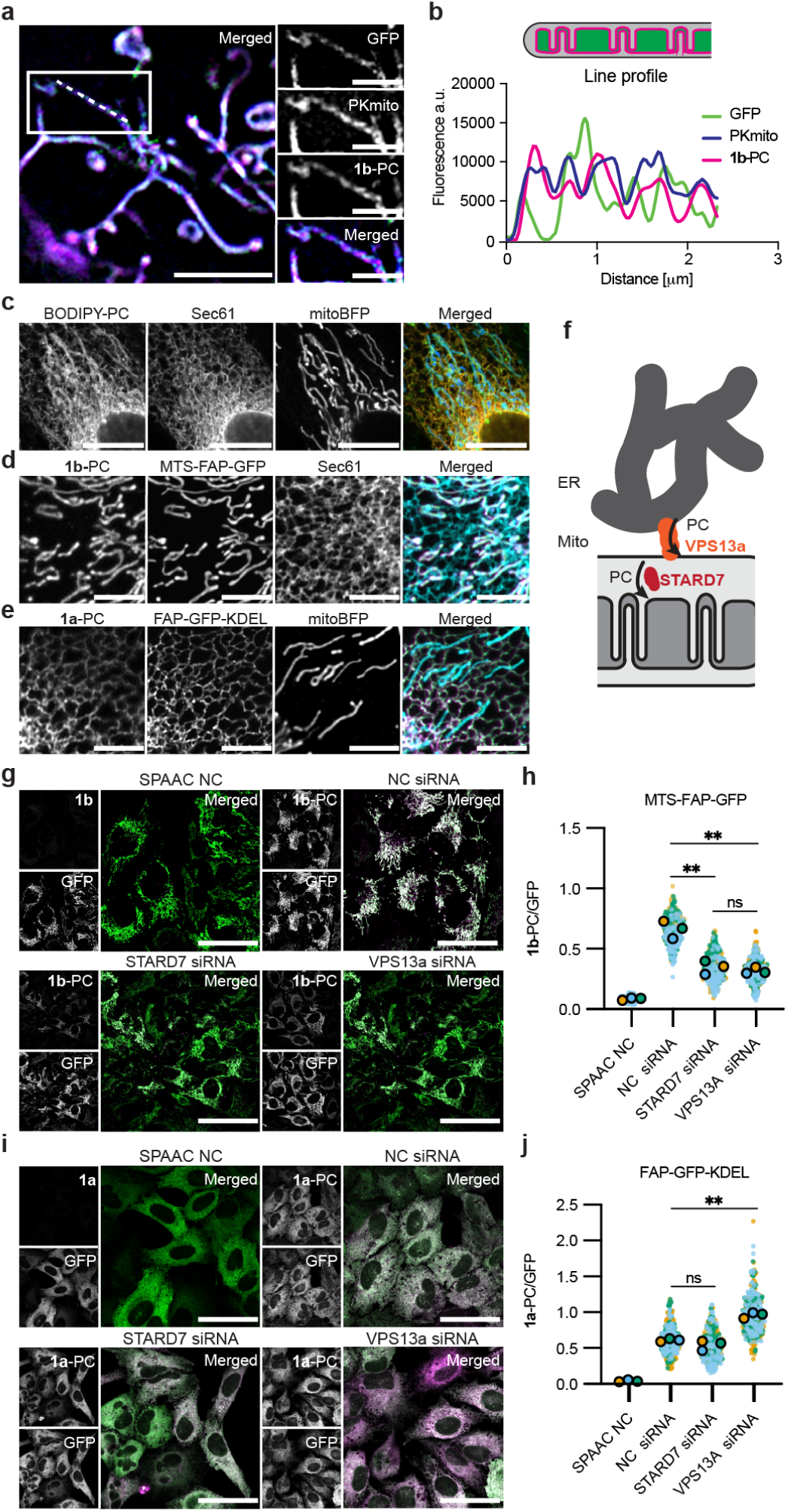
Using FACES to analyze PC trafficking between the mitochondrion and the ER. **a**, SIM^2^ of **1b**-PC detection in mitochondria of MTS-FAP-GFP cells counterstained with the cristae marker PKmito. Boxed region is magnified to the right. White line indicates the segment used for plotting the fluorescent intensity profile. Scale = 4 μm and 2 μm (inset). **1b**-PC fluorogenic signal forms a banded pattern that co-localizes with cristae. Additional example shown in Supp. Fig. 5d. **b**, Fluorescence intensity line profile from (**a**). **c,** Airyscan images show total N_3_-PC pools labeled with BODIPY-DBCO overlap with both the ER (sec61mCherry) and mitochondria (mitoBFP). 405, 488, and 561 nm lasers were used for excitation. Scale = 10 μm. **d,** Airyscan images of a HeLa cell expressing MTS-FAP-GFP and Sec61b-mCherry (cyan) show that **1b**-PC fluorescence detected in mitochondria does not overlap with the ER. 488, 561, and 631 nm lasers were used for excitation. Scale = 5 μm. **e,** Airyscan images of a HeLa cell expressing FAP-GFP-KDEL and mito-BFP (cyan) show that **1a**-PC fluorescence detected in the ER does not overlap with mitochondria. 405, 488, and 631 nm lasers were used for excitation. Scale = 10 μm. For experiments (**a**-**e**) 100 μM N_3_-Cho was incorporated for 24 h and 500 nM of either **1a** (ER) or **1b** (Mito) were used for PC labeling. **f**, Cartoon schematic depicting two proposed modes of PC transport into mitochondria. **g**-**h**, Knockdown of two putative ER-mitochondrial lipid transporters alters the distribution of PC between the two organelles. **g** and **i**, Representative LSM images of siRNA transfected HeLa cells stably expressing either MTS-FAP-GFP (**g**) or FAP-GFP-KDEL (**i**). 488 and 631 nm lasers were used for excitation. Scale = 50 μm. **e** and **g**, Quantified fluorescence intensity ratios of fluorogen (**1a** or **1b**) to GFP. Cells were treated with 25 pmol siRNA and 100 μM N_3_-Cho for 48 h prior to labeling with 500 nM **1a** (ER) or **1b** (Mito). SPAAC NC represent cells that were treated with NC siRNA but were not any fed N_3_-Cho. Experiments were conducted in triplicate; *n* = 3, *N* = 100-116 cells per condition. Statistics reflected paired two-tailed T-test of the population means.

### Lipid transporters contributing to PC trafficking between ER and mitochondria

PC is the most abundant lipid in mammalian cells and is a component of all organelle membranes, which make disentangling its subcellular transport mechanisms challenging. For instance, N_3_-PC pools strongly overlap between the ER and mitochondria when labeled with a constitutively fluorescent dye like BODIPY (BODIPY-DBCO) (Fig. 2c). However, using FACES, organelle specific N_3_-PC pools can be selectively illuminated and imaged. Cells expressing MTS-FAP-GFP detect on **1b**-PC in mitochondria that does not overlap with the ER marker Sec61b-mCherry (Fig. 2d). Likewise, cells expressing FAP-GFP-KDEL detect **1a**-PC in the ER that does not overlap with the mitochondria (Fig. 2e).

Utilizing the ability to quantify PC in mitochondria and ER, we tested how defects in lipid transport differentially affect these pools. For this, we generated stable cell lines expressing either FAP-GFP-KDEL (ER) or MTS-FAP-GFP (mitochondria) for further experiments. PC is synthesized in the ER and is transported to mitochondria by lipid transport proteins (LTPs)^22,25^. We used siRNA to silence two LTPs: VPS13a, a bridge-type LTP that interacts with VAPB at ER-mitochondria contact sites and transports bulk phospholipids^26^, and STARD7, a small shuttle-type LTP that localizes to the mitochondrial intermembrane space and is thought to transport PC between outer and inner membranes^27–29^ (Fig. 2f). Silencing resulted in a >50% reduction in both VPS13a and STARD7 protein content in each experiment (Supplementary Fig. 6). In both cases VPS13a and STARD7 siRNAs reduced **1b**-PC fluorescence in mitochondria by approximately 50% relative to negative control (NC) siRNA treated cells (Fig. 2g, h). However, only knockdown of VPS13a increased **1a**-PC fluorescence in the ER (Fig. 2i, j). This is consistent with the proposed role of VPS13A as an ER to mitochondrial LTP and that of STARD7 as an intra-mitochondrial LTP.

### Transbilayer asymmetry of multiple phospholipids generated at the trans-Golgi network

There currently exist few, if any, tools capable of measuring membrane asymmetry in living cells. We asked if the spatial specificity of FACES could provide this capacity when FAPs are fused to the transmembrane domain for leaflet-specific fluorogen detection. A potential site of asymmetry generation in the secretory pathway is at the TGN, where vesicles bud off from the Golgi for trafficking of proteins and lipids to the highly asymmetric PM. We explored asymmetric lipid detection in the TGN by reciprocal tagging of the transmembrane protein TGN38, at N- and C-termini, to selectively orient FAPs toward either the exoplasmic or cytoplasmic facing leaflets (Fig. 3a). Here only the transmembrane domain and short cytosolic tail of TGN38 were used, in conjunction with a flexible glycine-serine linker, in order to bring the FAP into close proximity to the membrane surface. Using this strategy we generated stable HeLa cell lines expressing either FAP-TGN38-GFP (exoplasmic leaflet) or GFP-TGN38-FAP (cytoplasmic leaflet) under the control of a doxycycline inducible promoter. TGN38 cycles to and from the PM via secretory vesicles and early endosomes^30^, thereby labeling a split population of trans-Golgi cisternae and trans-Golgi network vesiculated compartments^31^.

**Figure 3.**
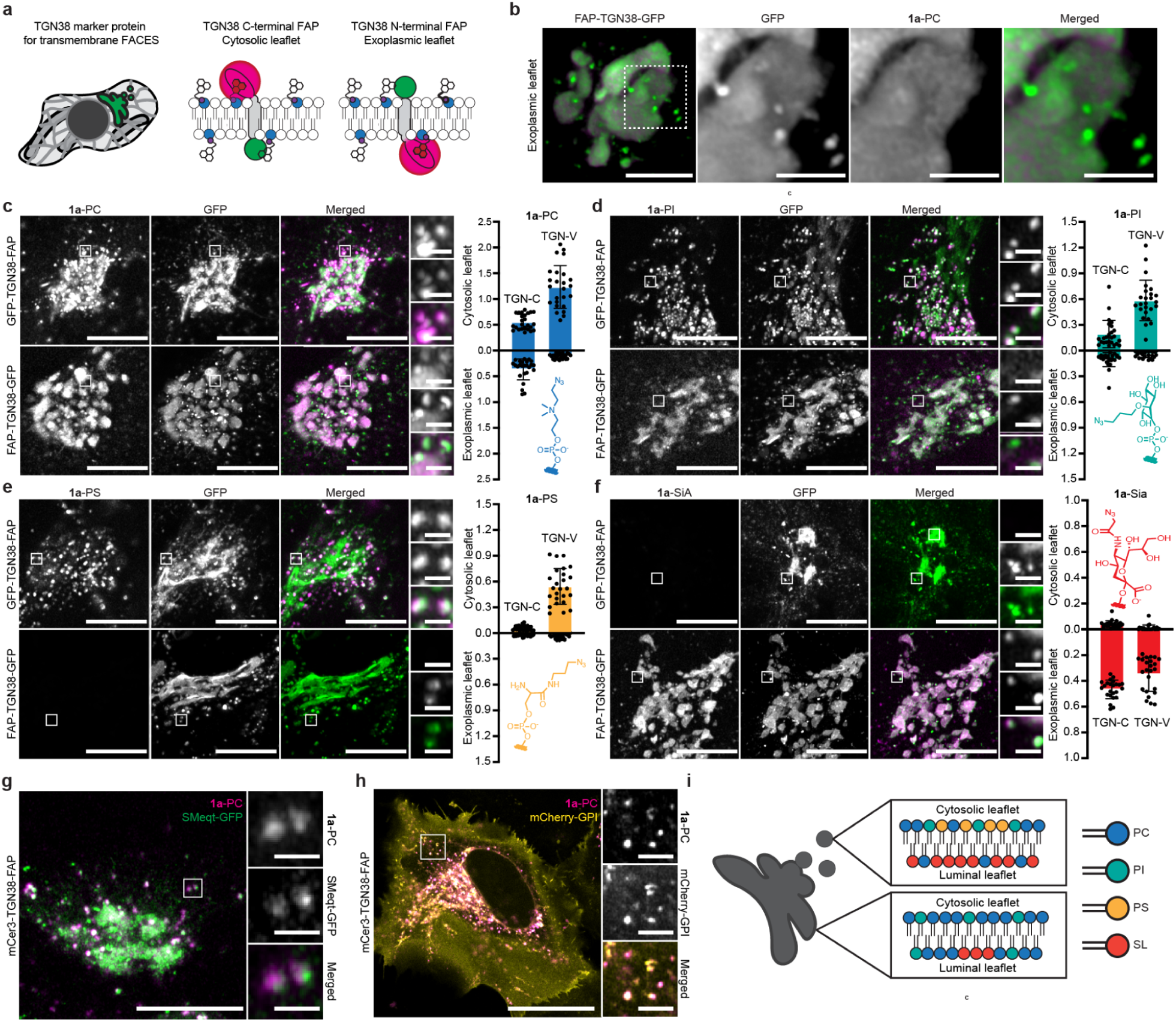
Imaging lipid asymmetry generated at the TGN with FACES **a,** Illustrated schematic representing the constructs used for reciprocal tagging of TGN38 with a FAP exposed to either the cytosolic or exoplasmic leaflet. **b,** 3D rendering of z-stacked Airyscan images reveal a lack of **1a**-PC labeling on the exoplasmic leaflet of vesicles connected to TGN cisternae. Magnified images from the boxed region are shown to the right. Scale = 5 μm and 2 μm (inset). **c**-**f,** Fluorescence intensity ratio of **1a** to GFP in TGN cisternae (TGN-C) and TGN vesicles (TGN-V) of the cytoplasmic leaflet (top) and exoplasmic leaflet (bottom) measured using LSM. Bar graphs represent mean population values and error bars reflect SD; *n* = 1, *N* = 25 cells. Example Airyscan images are provided as z-stacked maximum projections. Magnified insets show labeling of the TGN-V population. Structures of N_3_-containing lipid head groups are provided. Scale = 10 μm and 1 μm (inset). **g,** Colocalization of the secreted SM probe SMeqt-GFP and cytoplasmic **1a**-PC in individual TGN vesicles. Here, 488 and 633 nm laser lines were used for imaging to avoid imaging of mCer3. Scale = 10 μm and 1 μm (inset). **h,** Colocalization of mCherry-GPI and cytoplasmic **1a**-PC. Scale = 25 μm and 3 μm (inset). **i**, Conceptual model representing the proposed asymmetric distribution of PC and SM lipids that is generated alongside TGN vesicle formation.

Since these chimeric constructs lack the native N-terminal domain of TGN38, and could differ based on the reciprocal tagging strategy, it was necessary to investigate the identity of the vesicular population labeled by each construct. The fluorescent protein mCherry, with an N-terminal secretion signal peptide (SP-mCherry), was used as a marker for soluble secretory cargo and transfected in each cell line. Nearly all GFP-positive TGN vesicles colocalized with the mCherry marker in both cell lines (Supplemental Fig. 7a). Time lapse imaging further revealed TGN vesicles containing soluble secretory cargo rapidly leaving TGN cisternae (Extended Data Fig. 1). Next we used RAB5-BFP and RAB11-BFP as markers for early endosomes and recycling endosomes, respectively. RAB11 did not colocalize with GFP in either cell line (Supplemental Fig. 7c). However, in both cell lines a small subpopulation of GFP-positive TGN vesicles colocalized with RAB5 (Supplemental Fig. 7c). This is in agreement with retrograde trafficking of endogenous TGN38, which travel directly from the PM to the TGN in early endosomes that bypass the recycling endosome compartment^32^.

Next we tested the ability of FAP-TGN38-GFP and GFP-TGN38-FAP to detect **1a**-PC in the TGN using FACES. Cells expressing FAP-TGN38-GFP, oriented toward the exoplasmic leaflet, generated **1a**-PC fluorescence in the lumen of TGN cisternae that was either absent, or strongly diminished, in the vesicular population. Using Zeiss Airyscan imaging, lateral segregation between exoplasmic **1a**-PC enriched and depleted regions could be observed at sites of vesicular budding from the TGN cisternae (Fig. 3b). In contrast, cells expressing GFP-TGN38-FAP, oriented towards the cytoplasmic leaflet, generated **1a**-PC fluorescence in the TGN cisternae with very strong signal arising from the vesicular population. These observations were quantified using CFLSM and FACES by measuring the ratiometric fluorescence of TGN cisternae and the vesicular population (Fig. 3c). **1a**-PC was equally distributed between both leaflets of TGN cisternae, with a slight enrichment on the cytoplasmic leaflet. However, the abundance shifted between these compartments, with a strong comparative enrichment in the cytoplasmic leaflet of TGN vesicles (Fig. 3c, Supplementary Fig. 8a). These results are consistent with previous measurements of PC in freeze-fractured yeast cells, which showed a cytoplasmic leaflet enrichment arising in the Golgi^33^, although we cannot rule out other mechanisms by which lumenal labeling might be reduced in TGN vesicles.

We also asked if FACES could measure the asymmetry of non-choline lipids at the TGN. Recently, metabolic incorporation of azido-containing L-serine^11^ (C-L-Ser-N_3_) and *myo*-inositol^12^ (Ins-2-N_3_) have been shown to label PS and PI lipid classes, respectively. The latter is likely to label phosphorylated PIs^12^, albeit at lower abundances. Both PS and PI classes are synthesized in the ER and have been proposed to be trafficked to the cytoplasmic leaflet of the trans-Golgi and TGN through lipid transfer proteins^34,35^. Metabolic incorporation of Ins-2-N_3_ and C-L-Ser-N_3_ probes into N_3_-PI and N_3_-PS phospholipids was confirmed by high resolution LC-MS/MS (Supplemental Fig. 9). When analyzed with FACES, we observed **1a**-PI in Golgi cisternae that was evenly distributed between exoplasmic and cytoplasmic leaflets (Fig. 3d, Supplementary Fig. 8b). In TGN vesicles, the distribution of **1a**-PI shifted to the cytoplasmic leaflet, similar to that for PC. For **1a**-PS, using C-L-Ser-N_3_, there was an absence of signal on both leaflets of Golgi cisternae and a high, exclusively cytoplasmic abundance on TGN vesicles (Fig. 3e, Supplementary Fig. 8c). This is consistent with previous observations that the PS-binding C2 domain of lactadherin, when expressed in the cytoplasm, robustly labels TGN vesicles, but not Golgi cisternae^36^. As a control for these cytosolic leaflet-enriched phospholipids, we imaged the distribution of surface glycans in cells fed with peracetylated *N*-azidoacetylmannosamine (Ac_4_ManNAz)^37^, which is metabolized into sialic acid (Sia) within the cell. We measured an exclusively exoplasmic distribution of **1a**-Sia in both cisternae and vesicles (Fig. 3f, Supplementary Fig. 8d), consistent with the synthesis of protein and lipid surface glycans in the Golgi lumen^38^. Lastly, co-staining cells expressing mCer3-TGN38-FAP (Blue FP variant) with equimolar amounts **1a** and BODIPY-DBCO, a conventional fluorophore, demonstrated that FACES can spectrally separate organelle-specific lipid pools that overlap with total cellular phospholipid distributions (Supplementary Fig. 10)

Previous analyses of TGN vesicles^39^ and the PM^40^ suggest that loss of exoplasmic phospholipids could be substituted with SM, which is predominantly synthesized in the TGN and packaged in secretory vesicles. Consistent with this model, we observed that the SM probe SMeqt-GFP^17^ colocalized with vesicles enriched with cytoplasmic leaflet **1a**-PC (Fig. 3g) that rapidly exit the Golgi (Extended Data Fig. 2). SMeqt-GFP contains an N-terminal secretion signal peptide and passes through the secretory pathway, so it only binds the lumenal face of Golgi membranes. Similar results were obtained for vesicles labeled with the secretory cargo mCherry-GPI (Fig. 3h, Extended Data Fig. 3), which also co-traffics with SM^17^. Overall, these results support a model in which phospholipids become enriched on the cytoplasmic leaflet during TGN vesicularization, while the exoplasmic leaflet becomes enriched in newly synthesized sphingolipids in the TGN lumen (Fig. 3i). The resulting vesicles move both protein cargoes and asymmetrically-distributed lipids to and from the PM.

### PS asymmetry in the TGN is generated through lipid transport and flipping pathways

Genetic evidence in yeast suggests that the asymmetry of PS in the vesiculated TGN could be generated by P4-ATPase phospholipid flippases that translocate PS from the exoplasmic leaflet to cytoplasmic leaflet^41–45^. How asymmetry is generated in mammalian TGN has remained obscure and it is unknown to what extent lipid flipping, or transport pathways at ER-TGN contact sites, contribute to PS enrichment in the cytoplasmic leaflet (Fig. 4a). In mammalian cells, ORP10 and ORP11 have been characterized as PS transporters at ER-TGN contact sites that form PI-4-phosphate counter exchange complexes with ORP9 ^46–48^. Likewise, mammalian P4-ATPase phospholipid flippases play important roles in maintaining PS asymmetry in the PM and secretory pathway by redistributing lipids to the cytoplasmic leaflet^36,49–51^. To ensure that the C-L-Ser-N_3_ probe faithfully reports native PS distributions in the cells, we compared the localization of the PS-binding protein C2Lact-mCherry to that of N_3_-PS labeled with BODIPY-DBCO, which fluorescently labels intracellular N_3_-phospholipids. We found that BODIPY-conjugated N_3_-PS labeled the same intracellular membranes as C2Lact-mCherry (Fig. 4b).

**Figure 4.**
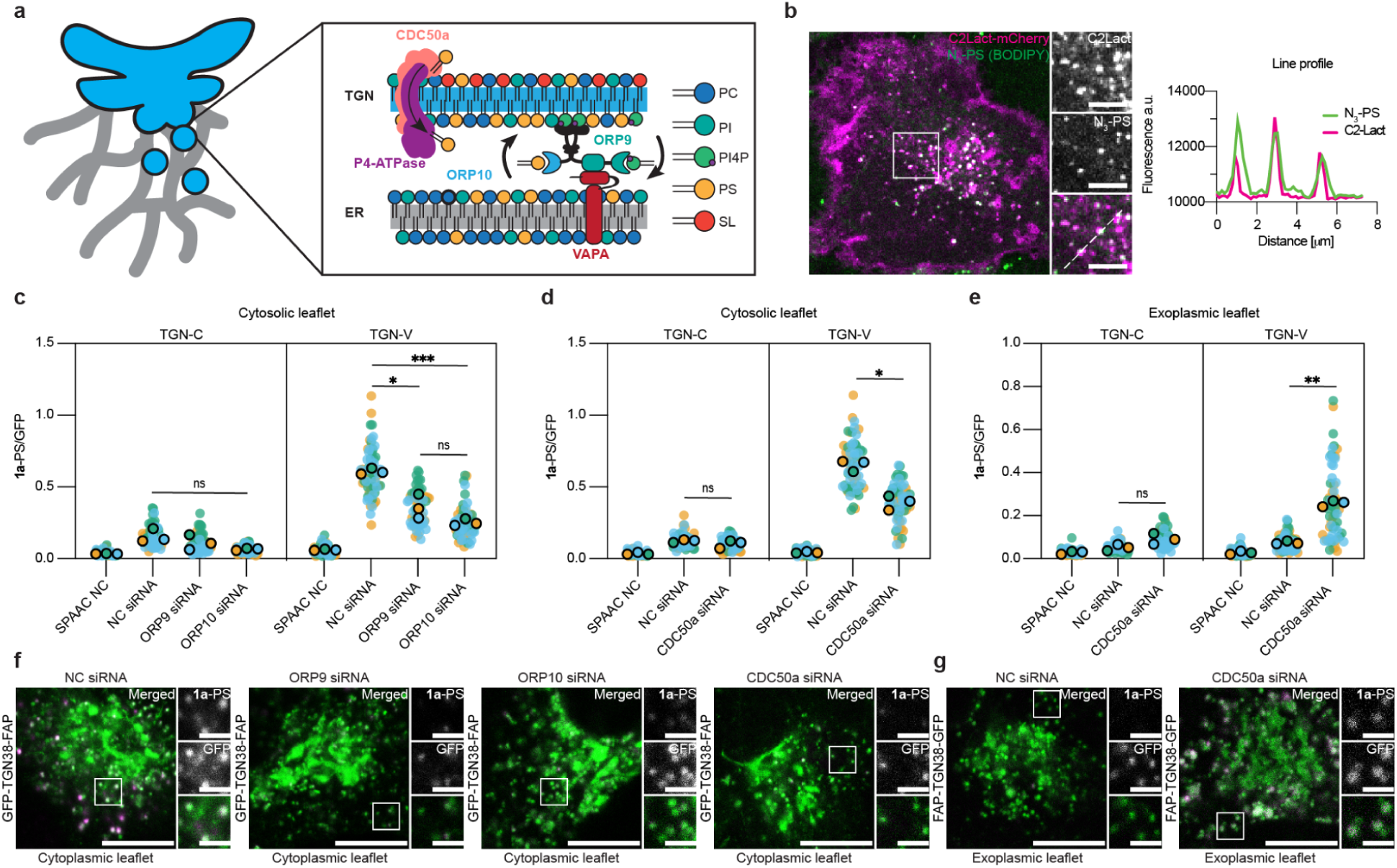
Mechanisms contributing to PS asymmetry at the TGN **a**, Cartoon schematic depicting two proposed lipid transport pathways generating PS asymmetry at the TGN. PS can be flipped from the exoplasmic leaflet to the cytoplasmic leaflet of the TGN by P4-ATPase/CDC50a flippases or transported from the cytoplasmic leaflet of the ER to the cytoplasmic leaflet of the TGN by ORP9/10 LTPs at contact sites with VAPA. **b,** Colocalization of the PS-binding probe C2Lact-mCherry with N_3_-PS lipids labeled with 1 μM BODIPY-DBCO. Magnified insets show colocalization of both markers in vesicles. Fluorescence line intensity profile from magnified inset is provided. Scale = 20 μm and 3 μm (inset). **c** and **d**, Quantified ratio of **1a**-PS to GFP fluorescence in the cytoplasmic leaflet of the TGN in response to siRNA transfection. **c**, ORP9 and ORP10 silencing reduce **1a**-PS relative to cells treated with a negative control (NC) siRNA. **d**, Silencing CDC50a also decreases **1a**-PS in the cytoplasmic leaflet. **e**, Silencing CDC50a increases **1a**-PS in the exoplasmic leaflet of the TGN-V relative to NC siRNA treated cells. Cells were incubated with 250 μM C-L-Ser-N_3_ and doxycycline (1 mg/mL) during the 48 h siRNA transfection period, followed by labeling with 500 nM **1a**. SPAAC NC represents cells that were not fed C-L-Ser-N_3_ as a staining control. In all experiments *n* = 3, *N* = 25 cells per condition; *, p < 0.05; **, p < 0.01; ***, p < 0.001, as determined by paired Student’s t-test. **f**, Representative LSM images from siRNA experiments plotted in (**c**) and (**d**) using the cytosolic leaflet reporter GFP-TGN38-FAP. **g**, Representative LSM images from siRNA experiments plotted in (**e**) using the exoplasmic leaflet reporter FAP-TGN38-GFP. Scale = 10 μm and 2 μm (inset). 488 and 633 nm lasers were used for illumination.

To gain mechanistic insight into how PS asymmetry is generated at the TGN, we used FACES to measure bilayer specific changes in N_3_-PS distribution in cells silenced for ORP9 and ORP10, or the essential P4A-ATPase subunit CDC50a^52,53^. Silencing of ORP9, ORP10, and CDC50a target genes was validated by western blot and resulted in >50% reduction in each protein (Supplementary Fig. 11a-d). Silencing either ORP9 or ORP10 reduced **1a**-PS by 39% and 58%, respectively, on the cytosolic leaflet of TGN vesicles compared to those treated with a non-targeted, negative control siRNA (Fig. 4c). In comparison, silencing CDC50a reduced **1a**-PS by 40% on the cytosolic leaflet of TGN vesicles (Fig. 4d). Some cells silenced for CDC50a were also positive for annexin V staining, indicating accumulation of extracellular PS in the PM (Supplementary Fig. 11e). We thus tested whether silencing CDC50a would also cause PS accumulation in the exoplasmic leaflet of the TGN lumen. Silencing CDC50a increased exoplasmic **1a**-PS content in vesicles (Fig. 4e). In these silencing experiments we did not notice obvious phenotypes in the TGN vesicle population despite the changes **1a**-PS (Fig. 4f, g).

Together these results highlight how different lipid transport mechanisms contribute to generating and maintaining PS asymmetry in the secretory pathway.

### FACES reports on changes to mitochondrial glycosylation dynamics

Based on our detection of surface glycans in the Golgi lumen, we asked whether FACES could be used as a general tool compatible for imaging of azido-sugars in living cells. Tetraacylated *N*-azidoacetylgalactosamine is widely used as a metabolic reporter for glycoconjugates containing *N*-acetylhexosamines (HexNAz)^54^. Ac_4_GalNAz fed to cells is converted to UDP-*N*-azidoacetylgalactosamine (UDP-GalNAz) by the hexosamine salvage pathway and is then further epimerized to UDP-*N*-azidoacetylglucosamine (UDP-GlcNAz) (Fig. 5a). This results in robust labeling of *N-* and *O-* linked glycoproteins on the cell surface^54,55^ (Fig. 5b), secretory compartments where they are formed, and endosomal compartments where they are recycled. Meanwhile, intracellular protein glycosylation is modulated by nucleocytoplasmic and mitochondrial *O*-linked β-*N*-acetylglucosamines (*O-*GlcNAc), which are added to serine/threonine residues by *O*-GlcNAc transferase (OGT) and removed by *O-*GlcNAcase (OGA)^54^ (Fig. 5c). *O-*GlcNAc cycling has emerged as a regulator of metabolism, signaling, and immune evasion of cancer cells^56–58^ and mitochondrial motility in neurons^59^. However, intracellular glycosylations are not readily observed by standard bioorthogonal imaging approaches due to the much more extensive incorporation of Ac_4_GalNAz into surface glycans; very little signal from conjugated conventional fluorophores colocalizes with mitochondria, for example (Fig. 5d). Currently, detection of *O-*GlcNAcylation is performed in fixed cells or bulk extracts using *O-*GlcNAc sensitive antibodies^60^.

**Fig. 5.**
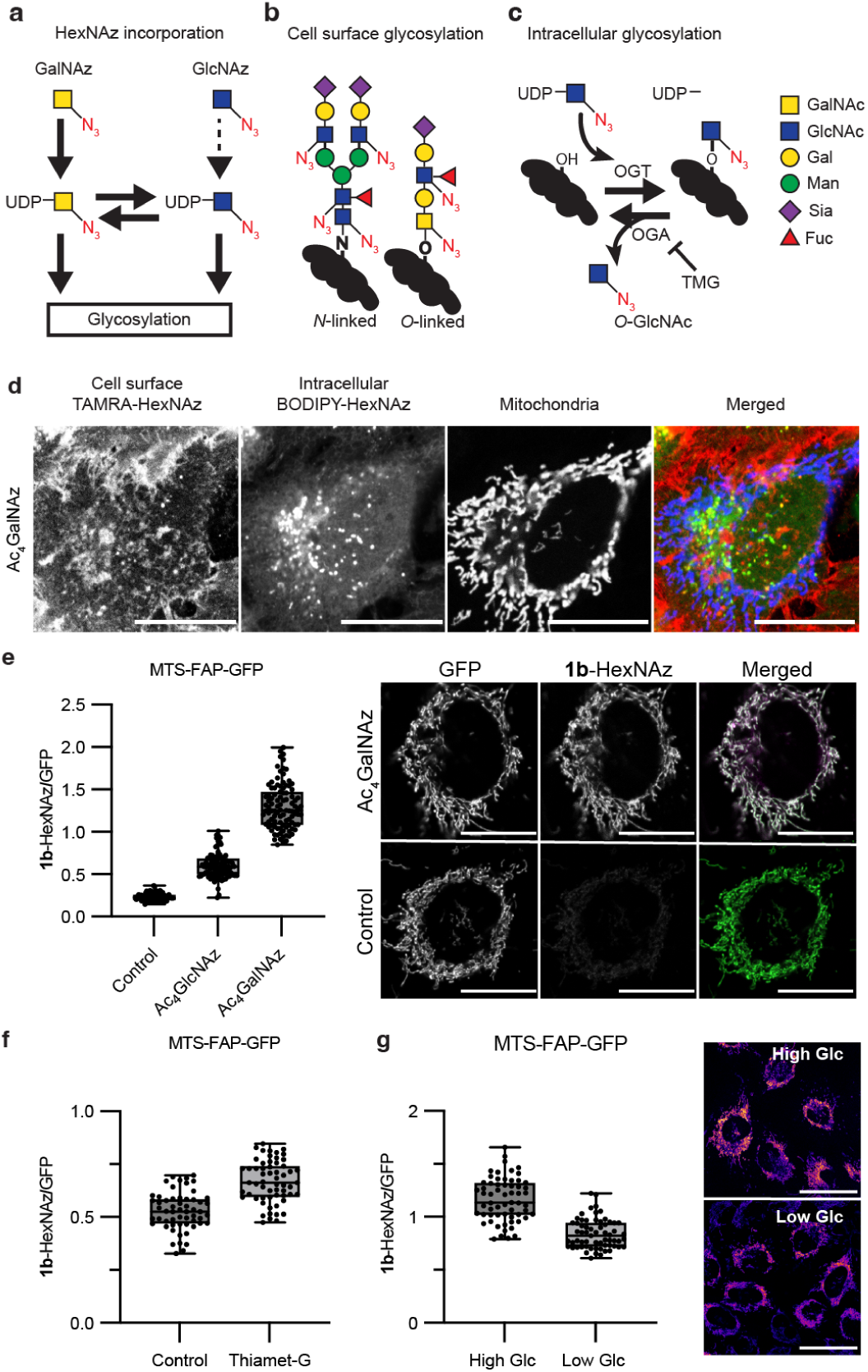
Intracellular imaging of *N-*acetylhexosamine sugars with FACES **a**, Illustration depicting the metabolic fate of Ac_4_GalNAz (yellow square) and Ac_4_GlcNAz (blue square) incorporation into cellular glycoconjugates containing *N*-acetylhexosamine sugars. Ac_4_GalNAz fed to cells is converted to UDP-GalNAz in the cytosol and further epimerized to UDP-GlcNAz, thereby robustly labeling both UDP-HexNAz pools (UDP-GalNAz + UDP-GlcNAz). Conversely, Ac_4_GlcNAz is not efficiently recycled to UDP-GlcNAz. **b**, UDP-HexNAz is incorporated into diverse extracellular glycoproteins proteins on the cell surface. Structures for an *N*-linked glycoprotein and *O*-linked mucin are shown as examples. **c**, UDP-HexNAz is incorporated into intracellular *O*-GlcNAcylated proteins. Protein glycosylation is catalyzed by *O*-GlcNAc Transferase (OGT) and can be reversed by *O*-GlcNAcase (OGA). OGA is pharmacologically inhibited by Thiamet-G (TMG). **d,** Bioorthogonal imaging of HeLa cells fed 100 μM GalNAz for 24h and labeled with 5 μM TAMRA-DBCO (cell surface) and 5 μM BODIPY-DBCO (intracellular), relative to the mitochondrial marker Mito-BFP. Scale = 20 μm. **e-i**, Stable MTS-FAP-GFP cells detect mitochondrial HexNAz. **e**, Quantified mitochondrial HexNAz incorporation in cells fed either 100 μM Ac_4_GalNAz or 100 μM Ac_4_GlcNAz for 24h, versus unfed controls; *n* = 1, *N* = 100 cells. Cells were labeled with 500 nM **1b** and washed as previously described. Representative images of Ac_4_GalNAz fed cells and control are shown on the right; scale = 20 μM. **f**, Response of mitochondrial HexNAz to OGA inhibition using Thiamet-G (TMG). Cells were treated with 50 μM Thiamet-G (or DMSO control) and fed 100 μM Ac_4_GalNAz for 24h. Displayed values are normalized to the average background fluorescence of unfed cells, which were used to control for **1b** labeling; *n* = 1, *N* = 59 cells. **g**, Response of mitochondrial HexNAz to glucose availability. Cells were fed 100 μM Ac_4_GalNAz and grown in DMEM containing either low (5.5 mM) or high (30 mM) glucose for 24h. Displayed values are normalized to unfed cells used as control for **1b** labeling; *n* = 1, *N* = 63 cells. Representative LSM images (right) show **1b**-HexNAz fluorescent intensities displayed using the FireLUT. Scale = 50 μm. Box plots reflect the first, median, and third quartiles, while whiskers reflect min to max values.

Using MTS-FAP-GFP expressing cells, we tested the efficacy of FACES in reporting mitochondrial-specific changes in HexNAz. Cells were grown in the presence or absence of either 100 μM Ac_4_GalNAz or 100 μM tetraacylated *N-*azidoacetylglucosamine-tetraacylated (Ac_4_GlcNAz) for 24h. Ac_4_GlcNAz is not efficiently incorporated by the hexosamine salvage pathway^54^ and thus was used as a less metabolically active surrogate. SPAAC staining with **1b** generated fluorescence signal intensities 10-fold greater in cells fed 100 μM Ac_4_GalNAz compared to unfed control cells (Fig. 5e). In contrast, **1b** fluorescence from cells fed Ac_4_GlcNAz, which is less efficiently incorporated into glycosylated proteins, was significantly lower than that for Ac_4_GalNAz. This indicates that the FACES signal from **1b**-HexNAz is from sugars that have been metabolically incorporated and not due to free azido sugars.

Next we tested whether FACES can report changes to mitochondrial *O-*GlcNAc levels resulting from pharmacological treatment or nutritional state. We found that the OGA inhibitor Thiemet-G increased **1b**-HexNAz fluorescence, suggesting that the fluorogen signal was responsive to the activity of *O*-GlcNAc modifying enzymes (Fig. 5f). In cancer cells, glucose uptake is positively correlated with *O-*GlcNAc levels, which in-turn reprogram cellular metabolism tailored towards glucose availability^61^. We observed that switching cells from high (30 mM) to low glucose (5.5 mM) levels during Ac_4_GalNAz labeling reduced **1b**-HexNAz fluorescence by 30% (Fig. 5g). This is similar to glucose-dependent changes in mitochondrial *O*-GlcNAc levels previously reported in breast cancer cells^62^. Taken together, these data demonstrate that FACES can report changes in mitochondrial HexNAz incorporation as a function of cell metabolism and *O*-GlcNAcylation in living cells, and support the hypothesis that mitochondrial *O-*GlcNAcylation is a post-translational modification that is responsive to the cell’s dietary state.

## Discussion

Studying cellular roles for non-proteinaceous biomolecules has long been hindered by the lack of robust tools for their imaging in live cells. Here we introduce a new approach, FACES, that combines the chemical specificity of metabolic incorporation with the spatial control of genetic protein tagging for imaging of cellular metabolites. FACES harnesses reversible fluorogenic systems, where fluorescence increases many orders of magnitude upon specific binding to a FAP. It combines this turn-on fluorescence with the specificity of bioorthogonal reactions for molecular sensing. This approach is powerful for the subcellular imaging of phospholipids, their transport between organelles, and orientations across membrane leaflets, but can also be applied to other biomolecules. Metabolic labeling has emerged as a particularly important tool for glycobiology and we demonstrate that FACES is responsive to labeled glycoproteins in mitochondria, which have been proposed to act as regulators of metabolic function^63^. Future applications of FACES could be further expanded through increases in the repertoire of metabolic labeling, bioorthogonal chemistry, and the spectral plasticity of the fluorogen employed.

Compared to other bioorthogonal imaging modalities, FACES has several key advantages for mechanistic lipid membrane biology. For organelle-defined imaging, protein targeting sequences are more specific and generalizable than corresponding chemically-encoded localization tags. We show that this capacity can be applied to the quantitative characterization of lipid distribution phenotypes affected by trafficking networks across membrane contact sites. Beyond organelle-specific imaging, FAPs can also be fused to specific TMDs to read out bilayer asymmetry. Here, FACES provides effective discrimination beyond what is possible with direct fluorescence imaging, as the distance between two bilayer leaflets is less than 4 nm. Fluorogens show no fluorescence when unbound to their corresponding FAP, so quantitative detection of conjugate products occurs through the resulting fluorescence signal, not a change in its localization as for lipid binding proteins. Because of this capacity, FACES is the only method, to our knowledge, that allows for measurement of membrane asymmetry in live cells.

Applications of FACES could be further expanded through increases in the repertoire of metabolic labeling, bioorthogonal chemistry, and the spectral plasticity of the fluorogen employed. Conjugation of fluorogens to biomolecules is dependent on tagged metabolite incorporation, which are continuously expanding across biological systems^64^ and molecular classes^65^. One concern with any metabolic labeling approach is that tagged molecules might not fully mimic native compounds, which is especially relevant for lipids with minimal headgroups. For several azido lipids, previous studies, as well as the data presented here, suggest that compounds localize across a range of cells as expected based off of comparative analyses with other tools^10,11^, though it is challenging to preclude all differences between labeled and unlabeled species. An additional limitation common to bioorthogonal approaches is the time required for conjugation and washing out of unreacted fluorogens (1 h); these might make the analysis of rapid redistributions of the underlying labeled species, like those relevant for lipid signaling^66^, challenging. Tuning fluorogenic reagents for faster conjugations^67^ and washing could decrease the time for processing steps to capture dynamic distributions.

Our experiments demonstrate that FACES is an especially powerful tool for investigating cellular mechanisms by which lipids are transported between organelles and across individual membranes, an intense area of investigation. A plethora of shuttle and bridge-like LTPs that perform lipid exchange at contact sites^6^ and flippases/floppases/scramblases that act on bilayer asymmetry^68^ have been identified and biochemically tested. However, these transporters are characterized by overlapping affinities, apparent redundancies, and close interactions with lipid metabolic enzymes; how they act in concert to shape the membrane heterogeneity that defines eukaryotic cells is still unknown. Here, we identify transporters that contribute to the asymmetry of several labeled phospholipid classes at the TGN. The Golgi has long been proposed as a site of asymmetry generation, given the co-occurrence of secretory cargo sorting to the PM and sphingolipid synthesis^69^. We show that for one phospholipid class, PS, cytosolic LTPs and transbilayer flippases both contribute to the asymmetry observed in TGN vesicles. These secretory compartments are enriched in GPI-anchored secretory cargoes that associate with the exoplasmic leaflet of the PM, suggesting a mechanism by which asymmetry at the PM is generated earlier in the secretory pathway. Lipid asymmetry generated at the TGN could also be relevant for sorting membrane proteins based on the asymmetric TMD sequences that are required for their PM targeting^3,70^. As a tool that provides spatially-defined molecular quantitation, FACES could be broadly powerful for untangling how lipid trafficking, generation of membrane asymmetry, and transport of cargoes simultaneouslyoccurs throughout eukaryotic cells.

## Acknowledgements

Nicolas-Frédéric Lipp provided discussions and comments, Gulcin Perkunnaz provided discussions and reagents, Jennifer Santini and the UCSD Microscopy Core (NS047101, OD030505) provided assistance with microscopy, Chris Burd and Jennifer Lippincott-Schwartz provided plasmids. The work was supported by the National Institutes of Health (R35-GM142960 to I.B. and R35-GM141939 to N.K.D.), the Paul G. Allen Family Foundation, and the National Science Foundation (CHE-2310263 to M.D.B.). R.J.B. acknowledges Agencia Estatal de Investigación and the Ministerio de Ciencia e Innovación (RYC2020-030065-I). The global polar phospholipid analyses described in this work were performed at the Kansas Lipidomics Research Center Analytical Laboratory. Instrument acquisition and lipidomics method development were supported by the National Science Foundation (including support from the Major Research Instrumentation program; most recent award DBI-1726527), K-IDeA Networks of Biomedical Research Excellence (INBRE) of National Institute of Health (P20GM103418), USDA National Institute of Food and Agriculture (Hatch/Multi-State project 1013013), and Kansas State University.

## Author Contributions Statement

W.M.M. and I.B. conceived the project and designed all experiments. All experiments were conducted by W.M.M. unless otherwise noted. R.B. synthesized the malachite green probes. C.H.K. conducted LC-MS/MS experiments. E.W. and B.H. assisted W.M.M. in cloning constructs. J.L., C.A.F., and M.D.B. synthesized and provided the inositol and serine azido probes. C.J.O. assisted in SIM experiments. N.K.D. provided expertise in bioorthogonal chemistry and support to R.B. and C.H.K.

## Methods

### Synthesis of fluorogen reagents

Synthesis schematic of compounds **1a** and **1b** are shown in Supplementary Fig. 1 and synthetic protocols are detailed in the Supplementary Text. Products were validated by LC-MS and ^1^H/^13^C NMR.

### Generation of constructs

FAPs were cloned from pcDNA3.1-KozATG-dH6.2-2XG4S-actin^21^, pcDNA3.1-kappa-myc-dK-2xG4S-TMst^71^, and pcDNA3.1-kappa-myc-dL5-2XG4S-mCer3-KDEL^72^ (Addgene #73263, #145766, #73209). These FAP sequences and the pcDNA3.1 backbone were used for generating the FAP-GFP-KDEL, GFP-KDEL, and MTS-FAP-GFP constructs used in transfection experiments. The Ig Kappa secretion signal peptide was used in conjunction with a KDEL sequence for ER-targeted constructs, while the targeting sequence from OTC was used for MTS constructs. For stable expression, FAP-GFP-KDEL and MTS-FAP-GFP coding sequences were subcloned into the sleeping beauty expression vector SBbi-PUR^73^. FAPs targeted to the TGN used only the transmembrane domain and cytoplasmic tail of TGN38 and included an N-terminal secretion signal peptide from VSV-G. The linker sequences (SGGGGSFGLSGL) and (AAGLAHGSGSG) were used in between the FAP and TGN38 sequences, for N-terminal and C-terminal tagged constructs, respectively. Reciprocally tagged FAP-TGN38-GFP and GFP-TGN38-FAP constructs were assembled into the SBtet-Pur expression vector. The GPI-anchor marker protein was generated by subcloning a SBP-mCherry-GPI sequence, received from the Lippincott-Schwartz lab, into pcDNA3.1. The secreted mCherry construct was generated by removing the SBP and GPI sequences and adding a stop codon. The GFP-EqtSM plasmid was received as a gift from the Burd lab. RAB5-BFP (Addgene #49147) and RAB11-BFP (Addgene #79805) were received from Addgene. Additional Sec61mCherry and Mito-BFP plasmids were received from the Lippincott-Schwartz lab. All constructs were validated by whole plasmid sequencing.

### Cell culture

HeLa (ATCC CCL-2) cells were grown in Dulbecco’s Modified Eagle Medium (Gibco 11995-065) containing glucose (4.5 g/L), glutamine (4.5 g/L), sodium pyruvate (110 mg/L), phenol red, 10% FBS, and 1% PenStrep, at 37°C and 5% CO_2_ atmosphere. For microscopy, cells were seeded into 35 mm glass bottom dishes (MatTek) coated with fibronectin (7.5 µg/mL). For transient expression, cells were transfected with Lipofectamine 3000 in Opti-MEM media for 4 h, and changed into normal growth medium containing azido-containing bioorthogonal metabolite for 24 h prior to SPAAC labeling and imaging. Stable cell lines expressing FAPs were generated using the Sleeping Beauty transposon insertion system. After transfection, cells were grown under selection with puromycin (10 µg/mL) for 2 weeks until all showed fluorescence. Doxycycline (1 µg/mL) was used to induce expression in SBtet-FAP-TGN38-GFP, SBtet-GFP-TGN38-FAP, and SBtet-mCer3-TGN38-FAP cell lines (Fig. 3, Fig. 4, Supplementary Fig. 7, Supplementary Fig. 8, Extended Data Fig. 1, Extended Data Fig. 2, Extended Data Fig. 3). Cells were routinely tested for mycoplasma by MycoStrip (InvivoGen).

### Metabolic labeling

1-azidoethyl-choline (N_3_-Cho, Jena BioScience) and resuspended in PBS to stock concentration of 50 mM. Ac_4_ManNAz, Ac_4_GalNAz and Ac_4_GlcNAz (Click Chemistry Tools) were resuspended in DMSO to stock concentration of 100 mM. L-Serine analogue C-L-Ser-N_3_ and *myo*-Inositol analogue Ins-2-N_3_ were synthesized as previously reported, verified by ^1^H/^13^C NMR and high resolution mass spectrometry, and resuspended in PBS to a stock concentration of 50 mM and 100 mM, respectively. In all cases, metabolic incorporation occurred for 24 h, except for experiments in siRNA experiments Fig. 2 and Fig. 4, which occurred for 48 h. Afterwards, cells were washed once with HBSS buffer, and incubated in fresh growth media for 30 min to allow for metabolic consumption of any remaining bioorthogonal metabolites. For experiments with peracetylated sugars, 1 h incubation was used.

### SPAAC conjugation

Culture media was removed by aspiration and replaced with staining media for SPAAC labeling. The staining media contained 500 nM **1a** or **1b** diluted in serum-free DMEM clear (Gibco 31053-028), and incubated for 15-20 min at 37°C and 5% CO_2_ atmosphere. Cells were washed with 4 volumes of culture media containing 10% FBS, 3 times, at 15 min intervals to remove unreacted dye. For experiments with the mitochondrial-localized FAP, cells were incubated for an additional 45 min, followed by a fourth wash step prior to imaging. BDP-FL--DBCO was purchased from BroadPharm (BP-23473) and 1-2 µM was used for constitutive fluorescent labeling of azido-containing metabolites.

### Lipid analysis

Lipids were extracted with a modified Bligh and Dyer protocol. Briefly, cell pellets from T75 culture flasks were resuspended in 200 μL PBS and extracted with 2 mL 1:2 CHCl_3_:MeOH (v/v) and cell debris were removed by centrifugation. Extracts were transferred to new vials and 1 mL chloroform was added, followed by 1 mL H_2_O, and shaking. The lower organic phase was collected after centrifugation and the aqueous phase was extracted twice more with 650 μL and pooled. Lipid extracts were then dried under N_2_ and resuspended in 100 μL 1:2 CHCl_3_:MeOH for thin-layer chromatography (TLC) and 500 μL methanol for LC-MS/MS. For TLC, entire lipid extracts were spotted onto 10×10 cm HPTLC silica gel 60 plates, developed with 14:5:1 CHCl_3_:MeOH:NH_4_, and imaged with a Typhoon FLA 9500 using 633 nm laser. For high resolution LC-MS/MS (Supplementary Fig. 2a, 2b, 9), 5 μL sample was injected, and HRMS spectra of the total lipid extracts from HeLa cells were recorded on a Q Exactive Orbitrap mass spectrometer (Thermo Fisher Scientific) connected to a Vanquish Flex Analytical UHPLC System (Thermo Fisher Scientific) equipped with a C8 column (100 x 2.1 mm, particle size 1.9 μm). Following HPLC method was used for all runs using 0.1% formic acid in water (Solvent A) and 0.1% formic acid in methanol (Solvent B): 0-1 min 30% B, 1-4 min 30%-90% B, 4-15 min 90-99% B, 15-18 min 99%B, 18-20 min 30% B. Lipid standards (POPC, N_3_-DSPC) were supplied by Avanti Polar Lipids. Global polar phospholipid analysis (Supplementary Fig. 2c-d) was performed by ESI-MS/MS at the Kansas State Lipidomics Research Center as previously described^74–76^.

### Image collection

Confocal micrographs were acquired on a Zeiss LSM 880 microscope equipped with plan-apochromat 63x/1.4 NA or 20x/0.8 NA objectives. For far-red fluorogen-FAP signals, a 633 nm HeNe laser was used at 1-3% power. FP or other fluorophores were excited with 488 nm Argon (GFP), 405 nm (BFP/mCer3) or 561 nm (mCherry) diode lasers. All images used for intensity quantification were acquired in LSM mode using a QUASAR GaAsP detector using identical settings. The calculated power density for individual experiments are as follows: FAP-GFP-KDEL (Fig. 2g, h), 963 W/cm^2^ (488 nm) and 2600 W/cm^2^ (633 nm) with a pixel dwell time of 1.02 µs; MTS-FAP-GFP (Fig. 2i, j), 375 W/cm^2^ (488 nm) and 2600 W/cm^2^ (633 nm) with a pixel dwell time of 1.02 µs; GFP-TGN38-FAP and FAP-TGN38-GFP (Fig. 3c-d, Fig. 4c, d, f), 535 W/cm^2^ (488 nm) and 2600 W/cm^2^ (633 nm) with a pixel dwell time of 2.05 µs; FAP-TGN38-GFP (Fig. 4e, g), 1,017 W/cm^2^ (488 nm) and 3,800 W/cm^2^ (633 nm) with a pixel dwell time of 2.05 µs. Images highlighting fluorogen distribution (Figs. 1e, 2c, 2d, 2e, 3b, 3c, 3d, 3e, 3f, 3g, 3h,; Supplementary Figs. 7a, 7c, 10; Extended data Figs. 1, 2, 3) were acquired with an Airyscan detector and processed using default settings in Zen Black; these were not used for quantification due to non-linear processing. SIM imaging of mitochondrial **1b**-PC was performed on an Zeiss Elyra 7 Lattice-SIM^2^ system equipped with a plan-apochromat 100x/1.46 NA objective using 642 nm and 488 nm diode lasers (10% power each). Images were acquired in ZEN Black using Weak Live default settings with a pixel dwell time of 843 ms.

### Image quantification

All intensities were calculated using ImageJ on unprocessed images. For proof of concept experiments using transfected cells (Fig. 1), line profiles drawn were across 10 individual ER segments per cell and the total fluorescence ratio between fluorogen and GFP emissions were tabulated across them. For per-cell data (Fig. 1f, 1g, 1h), the 10 segments were averaged and 10 cells were analyzed per condition. Raw per segment fluorescence intensity is shown for all 100 segments (Fig. 1h). For mitochondrial and ER quantification (Fig. 2), fields of cells stably expressing to corresponding FAPs were analyzed by segmenting individual cells and computing the total fluorescence ratio between fluorogen and GFP emission across the entire cell, given the highly specific distribution of the signal for the compartment of interest. For TGN cisternae quantification (Fig. 3, Fig. 4), line segments were drawn across the Golgi apparatus of individual cells so that they did not intersect any TGN vesicles. For TGN vesicle quantification, 5 line segments were drawn across Golgi vesicles that did not intersect Golgi cisternae. For each compartment, the total fluorescence ratio between fluorogen and GFP were calculated across each and averaged together for a single cell value. Values were measured this way for 25 cells per condition (exoplasmic vs. cytosolic for PC, PS, PI, and SiA). Mitochondrial quantification in Fig. 5 was carried out identically as in Fig. 3. All statistical analyses were performed in GraphPad Prism 9.

### Gene knockdown

Silencer Select siRNAs were supplied by Thermo Fisher: STARD7 (s32370), VPS13a (s531256), ORP9 (s41693), ORP10 (s223235), CDC50a (s31427), and negative control 1 (4390843). Cells were transfected with Lipofectamine RNAiMAX (ThermoFisher) complexed with 25 pmol siRNA for 48h prior to analysis. 50 pmol siRNA was used for CDC50a. Total protein was extracted using RIPA Lysis and Extraction buffer (ThermoFisher) and measured by Bradford assay (Bio-Rad). A total 20 μg protein per extract was boiled in 1X reducing sample buffer and used for SDS-PAGE (BioRad, 4561083). Validation of gene knockdowns was performed by Western blotting using primary antibodies against STARD7 (Proteintech, 15689) 1:2500, VPS13a (Atlas antibodies, HPA021662) 1:1000, ORP9 (Proteintech, 11879-1-AP) 1:1,000, ORP10 (Proteintech, 15491-1-AP) 1:1000, CDC50a (Invitrogen, PA5-99762) 1:1000, α-Tubulin (ThermoFischer, 32-2500) 1:5000, and Actin (Invitrogen, PA5-78715) 1:5000. HRP-conjugated goat anti-rabbit IgG secondary antibody (Abcam, 6721; 1:10000) and goat anti-mouse secondary antibody (Invitrogen, PI31430; 1:10000) and Pierce ECL western blotting substrate (Thermo Fisher, 32209) were used for detection.

## Data Availability

All data are available in the Article or its Suppementary Information. Plasmids generated in this study have been deposited in Addgene.

## Supplementary Figures

**Supplementary Fig. 1.**
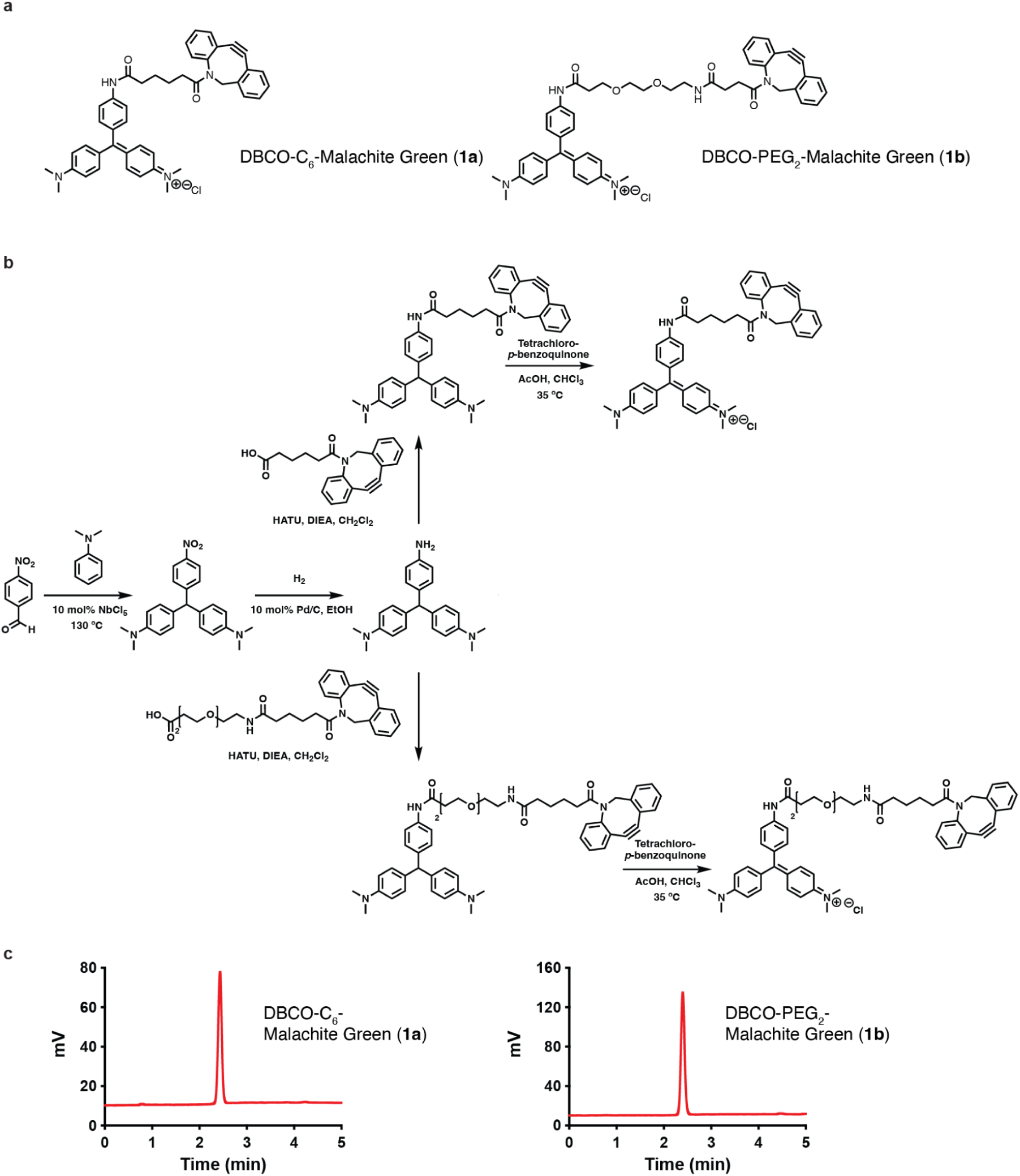
Synthesis of DBCO-linked fluorogens **a**, Chemical structures of all the Malachite Green (Chromatic forms) derivatives used in this study. **b**, Synthesis of the chromatic forms of DBCO-C_6_-Malachite Green (**1a**) and DBCO-PEG_2_-Malachite Green (**1b**). **c**, HPLC/ELSD spectrum corresponding to the chromatic forms of DBCO-C_6_-Malachite Green (**1a**) and DBCO-PEG_2_-Malachite Green (**1b**). Retention times were verified by mass spectrometry.

**Supplementary Fig. 2.**
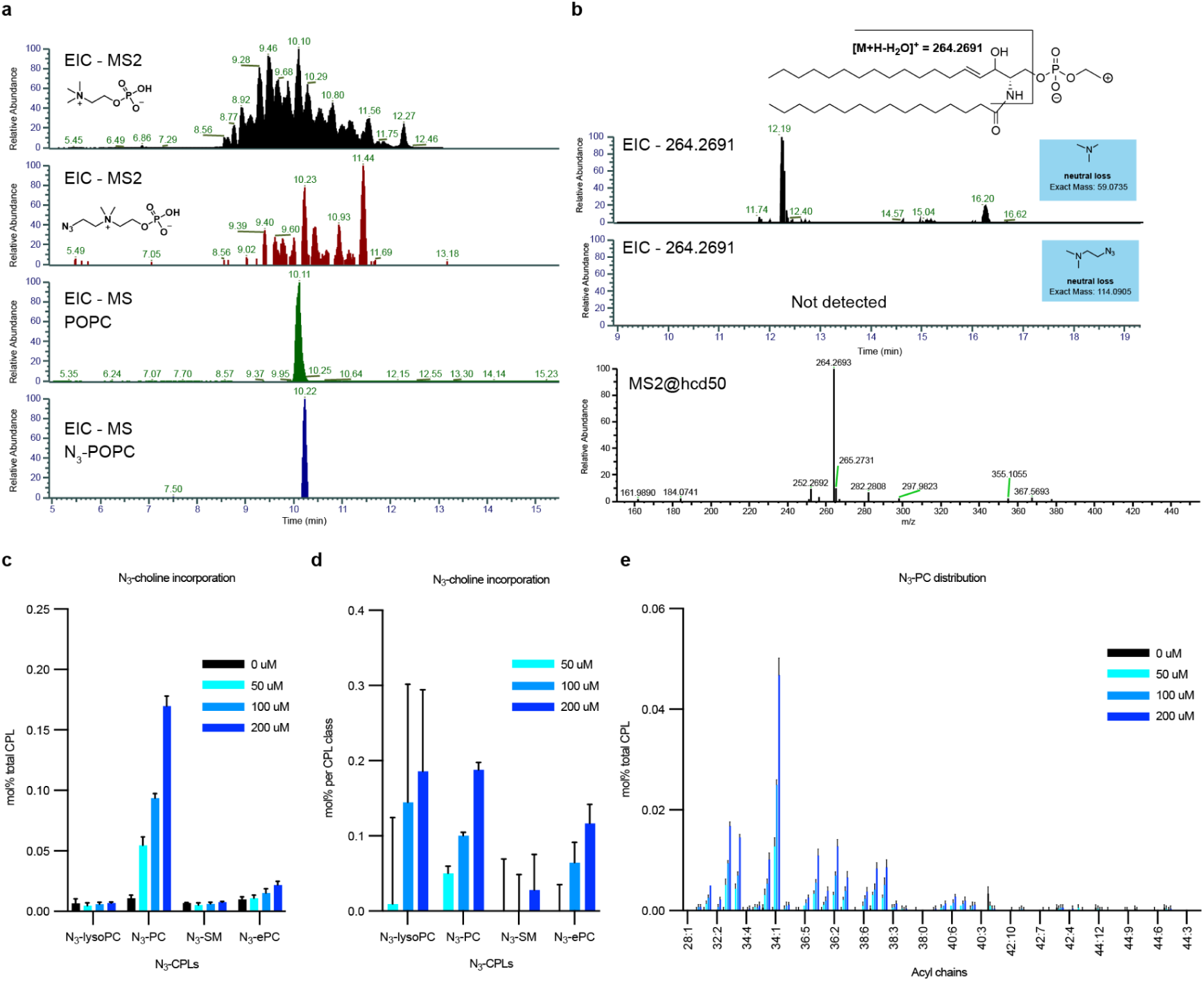
N_3_-Cho is metabolized into N_3_-PC but not N_3_-SM **a** and **b**, Cells were fed 100 μM N_3_-Cho for 24 h and extracted lipids were analyzed by LC-MS/MS. **a**, Comparative extracted ion chromatograms (EIC) from full scan/all ion fragmentation mode. MS2 fragments: choline (black), N_3_-Cho (red). EIC of POPC (green) and N_3_-POPC (blue) were detected as representative PC species in the sample. **b**, Sphingoid base fragment 264.269 was detected in neutral loss scan mode NL = 59 (choline containing lipids) but not in NL = 114 (N_3_-Cho containing lipids), indicating N_3_-Cho was not incorporated into SM in a detectable amount under these labeling conditions. **c**-**d**, Quantification of N_3_-Cho incorporation into choline phospholipids by ESI-MS/MS. **c,** Abundance of N_3_-Cho lipids represented as mol% relative to the total choline phospholipids in the cell. **d**, Accumulation of N_3_-Cho lipids represented as mol% relative to each lipid class. Minor levels of N_3_-SM is detected only at high labeling concentrations (200 μM N_3_-Cho) **e**, Distribution of N_3_-PC species by acyl chain length and saturation.

**Supplementary Fig. 3.**
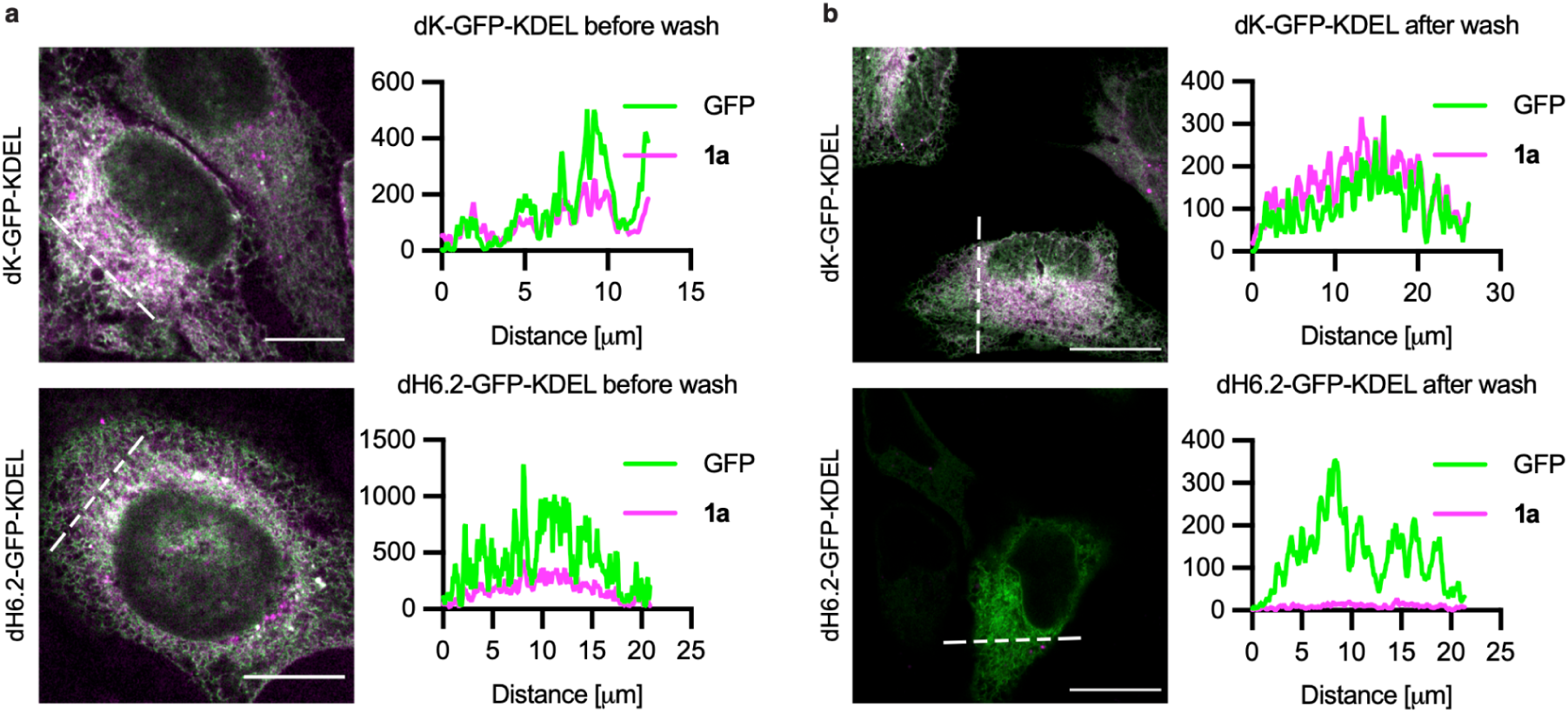
1a-FAP complex reversibility depends on the heavy chain scFvs FAP clone dH6.2 Cells that were not fed with any azido metabolites were transfected with either dH6.2-GFP-KDEL or light chain scFvs FAP variant dK-GFP-KDEL and stained with 500 nM **1a**. dH6.2 was shown to have reversible binding to MG^21,71^, while dK features nanomolar affinity^21,71^ and is thus irreversibly bound under experimental conditions. **a**, Cells imaged while in staining solution show colocalization of sensor (GFP) and fluorogen (**1a**) signal for both constructs. Scale = 10 μm. **b**, Cells imaged after washing 3x with DMEM show only fluorogen signal for the dK construct (top), as FAP-bound 1a is not washed out. Line profiles show the relative change in **1a** fluorescence intensity compared to GFP. Specific detection of SPAAC-conjugated fluorogens thus requires a lower affinity FAP like dH6.2. Scale = 25 μm.

**Supplementary Fig. 4.**
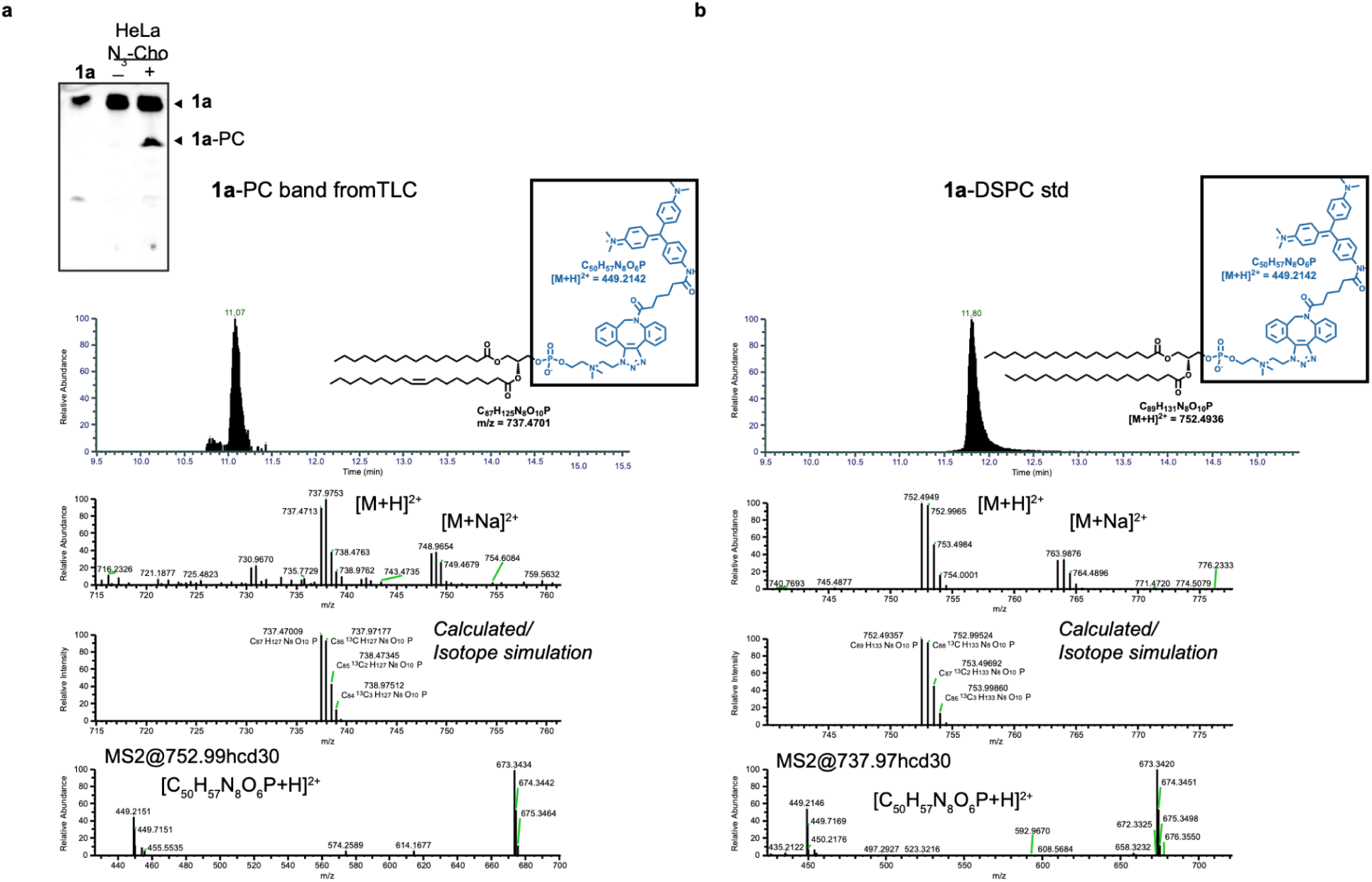
1a-PC conjugates are formed in cells **a,** HR-MS/MS analysis of **1a**-PC band from thin layer chromatography detecting [M+H]^2+^=747.4713 and MS2 fragment 449.2146. Cells stained with **1a** were not washed prior to lipid extraction to highlight conversion of **1a** to **1a**-PC. **b**, Comparative HR-MS/MS analysis of the **1a**-DSPC standard previously synthesized by reaction of **1a** with N_3_-DSPC detecting [M+H]^2+^=752.4949 and MS2 fragment 449.2146.

**Supplementary Fig. 5.**
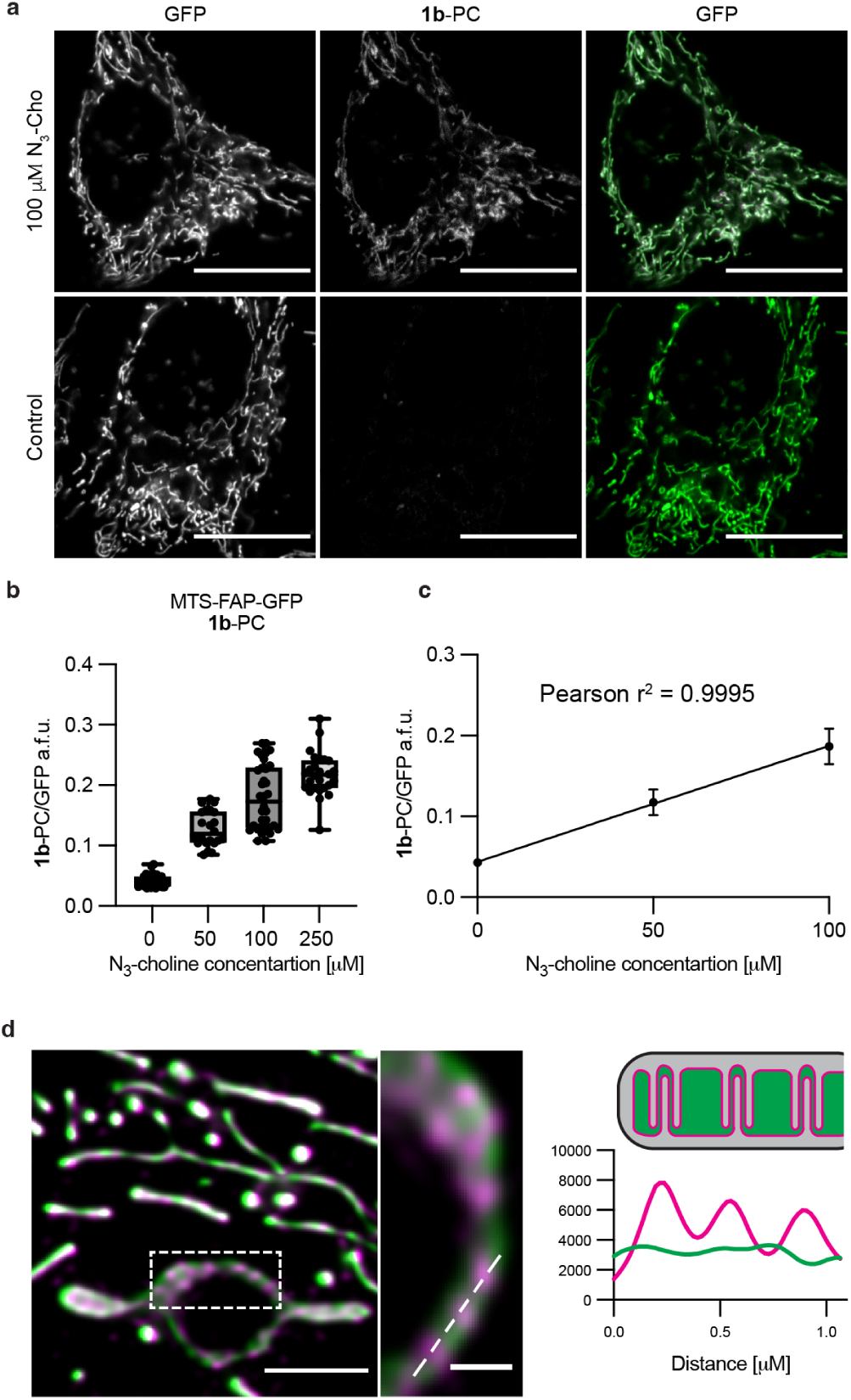
Linear quantification of 1b-PC in the mitochondrial matrix **a**, HeLa cells stably expressing MTS-FAP-GFP were fed different concentrations of N_3_-Cho for 24h and stained with 500 nM **1b**. Images show that fluorogen fluorescence (middle) is dependent on azido metabolic incorporation. Scale = 20 μm. **b**, Quantification of **1b**/GFP fluorescence in cells expressing MTS-FAP-GFP with increasing concentrations of N_3_-Cho; *N =* 24-34 cells. **c**, Linear increase of **1b**/GFP fluorescence only up to 100 μM N_3_-Cho. **d,** Super-resolution live cell imaging of **1b**-PC (magenta) in the mitochondrial matrix with Elyra 7 lattice SIM^2^. The graph below indicates fluorescence intensity profiles of **1b**-PC (magenta) and GFP (green) along the white dashed line (inset). Scale = 2 μm and 0.5 μm, respectively.

**Supplementary Fig. 6.**
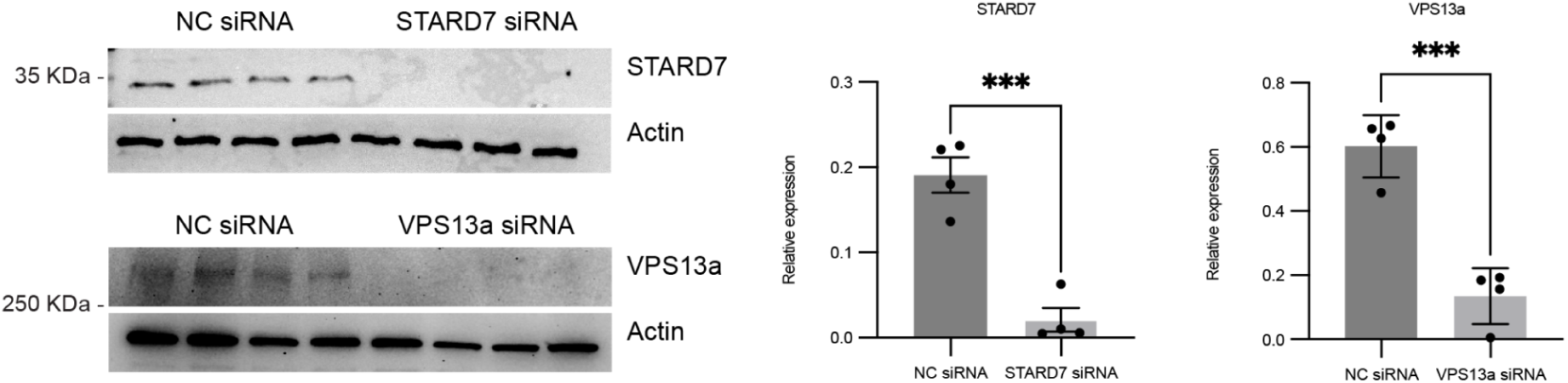
Silencing efficiency of STARD7 and VPS13a siRNAs Cells were transfected with 25 pmol siRNA for 48 h and silencing efficiency was determined by western blotting. Antibodies for STARD7 and VPS13a were used at 1:1000 dilution, while α-tubulin and actin was used as a loading control at 1:5000 dilution. Bar graphs reflect average expression relative to actin and error bars indicate SD (*n* = 4). Statistical significance was determined by Two-tailed Student’s T-test.

**Supplementary Fig. 7.**
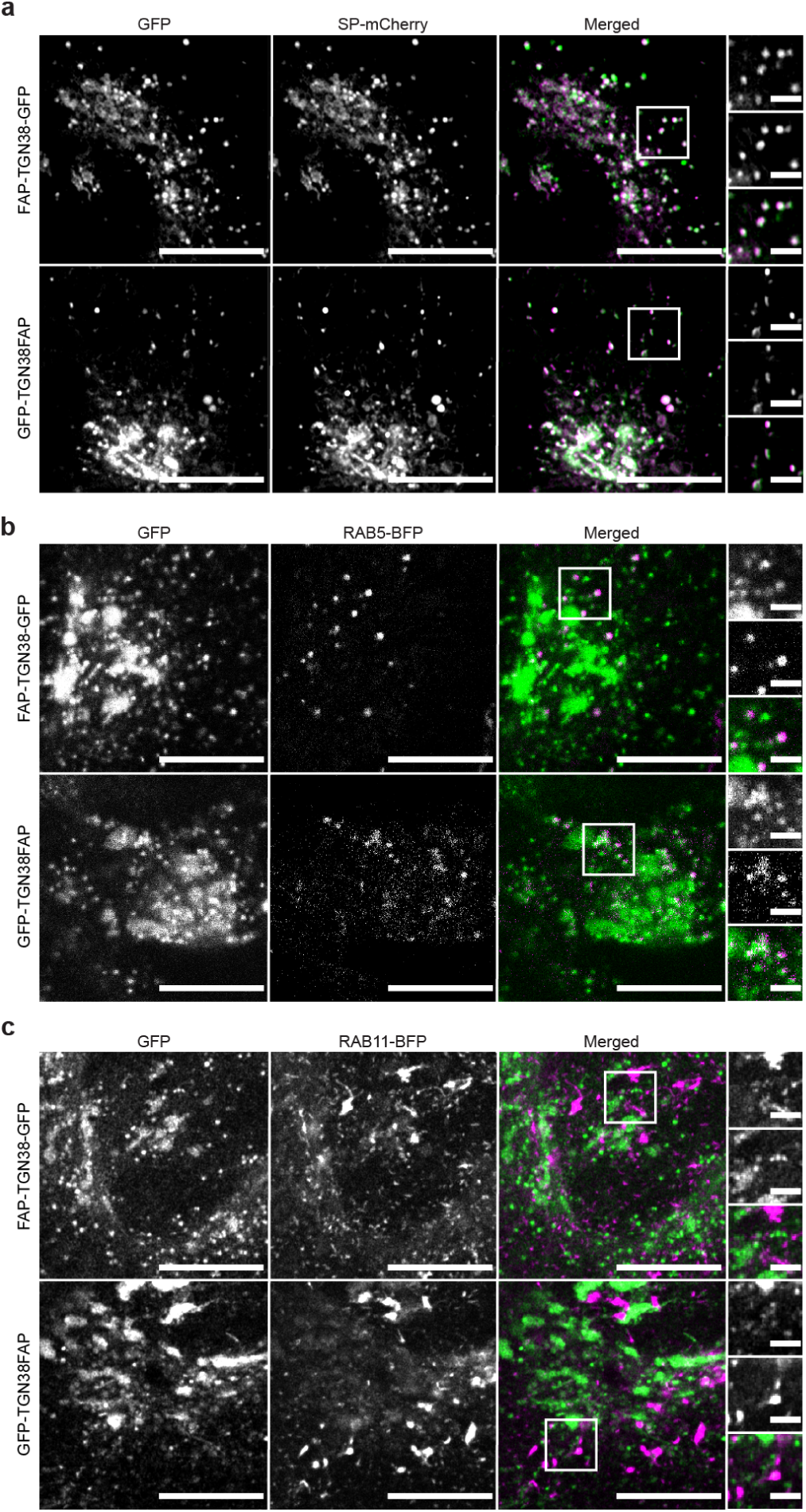
Colocalization between TGN constructs and markers for soluble secretory cargo and endosomes **a**, Airyscan images of cells expressing secreted mCherry (SP-mCherry; magenta) as a marker for soluble secretory cargo. SP-mCherry colocalized with a high proportion of TGN-Vs in both cell lines. **b**, LSM images of early endosomes labeled by RAB5-BFP (magenta) showed colocalization with a subset of TGN-Vs in both cell lines. **c**, Airyscan images of recycling endosomes labeled by RAB11-BFP (magenta) did not colocalize with TGN-Vs. Scale = 10 μm and 2 μm (inset). Images were acquired using 405, 488, and 561 nm lasers.

**Supplementary Fig. 8.**
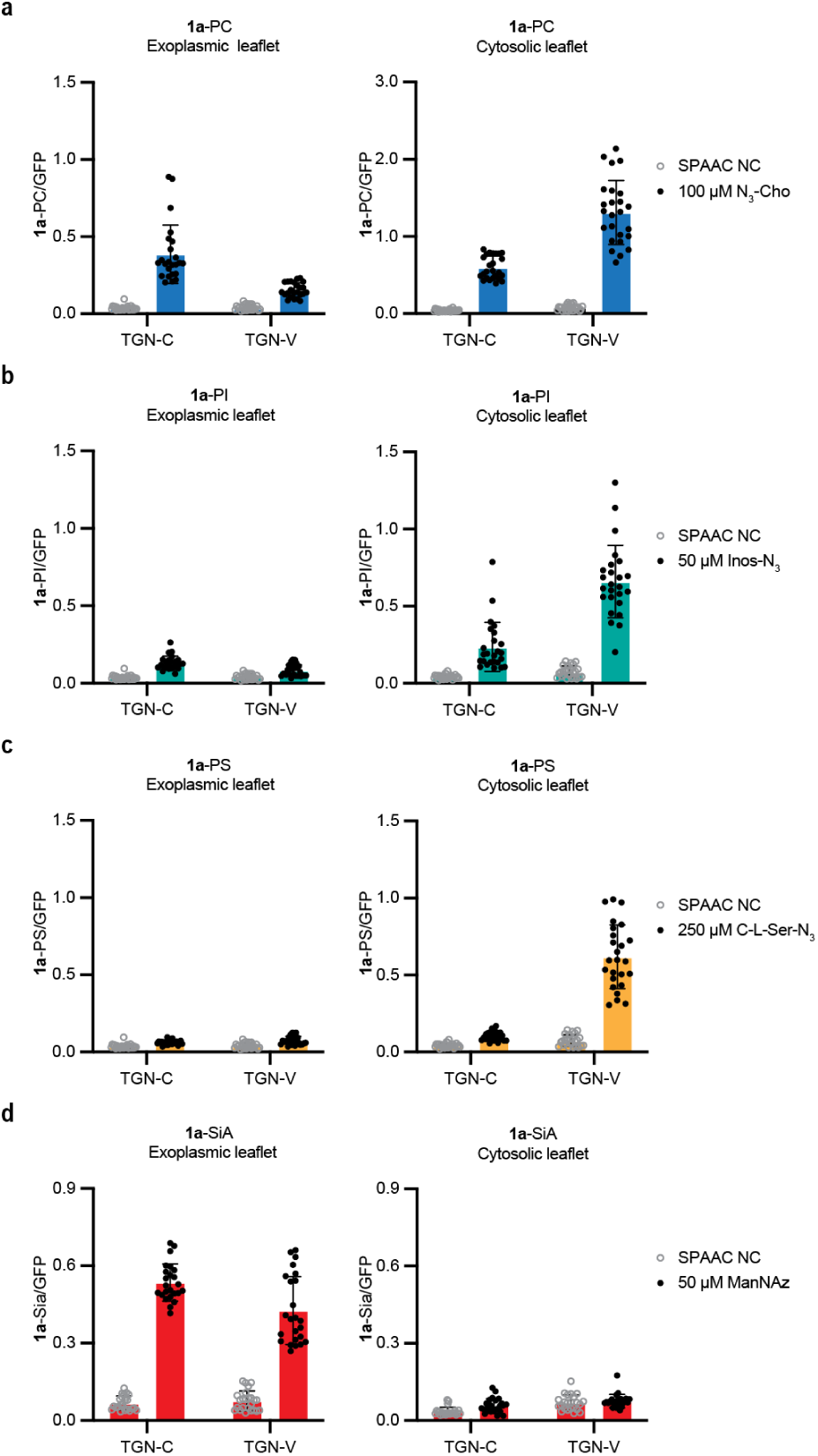
FACES quantification of phospholipid asymmetry across the TGN. Cells stably expressing doxycycline-inducible FAP-TGN38-GFP (lumenal leaflet) or GFP-TGN38-FAP (cytosolic leaflet) constructs were grown under 1 μg/mL doxycycline and fed different azido metabolic labeling reagents for 24h prior to staining with 500 nM **1a**. **a,** cells fed 100 μM N_3_-Cho. **b**, cells fed 50 μM Ins-2-N_3_. **c**, cells fed 250 μM C-L-Ser-N_3_. **d**, cells fed 50 μM Ac_4_ManNAz. LSM images were used for quantification. Structures of N_3_-containing lipid head groups are provided. In all experiments *N* = 25 cells per condition.

**Supplementary Fig. 9.**
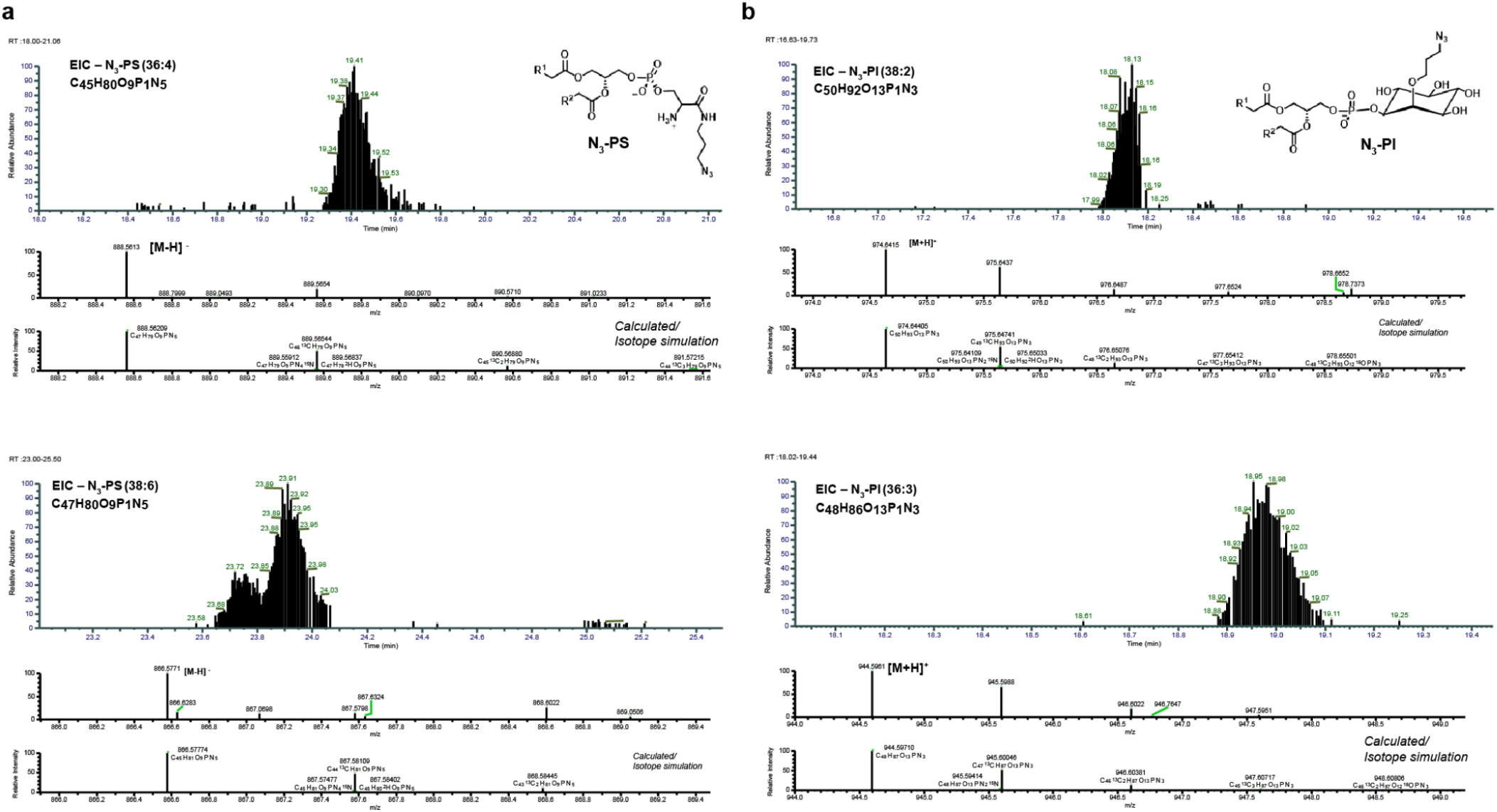
C-L-serine-N_3_ and Ins-2-N_3_ probes are metabolized into phospholipids **a**, Cells were fed 250 μM C-L-serine-N_3_ for 24 h and extracted lipids were analyzed by HR LC-MS. Extracted ion chromatograms (EIC) and full MS spectra of two representative N_3_-PS species 36:4 (top) and (38:6) (bottom) are given. **b**, Cells were fed 100 μM Ins-2-N_3_ for 24 h and extracted lipids were analyzed by HR LC-MS. EIC and full MS spectra of two representative N_3_-PI species 38:2 (top) and 36:3 (bottom) are shown.

**Supplementary Fig. 10.**
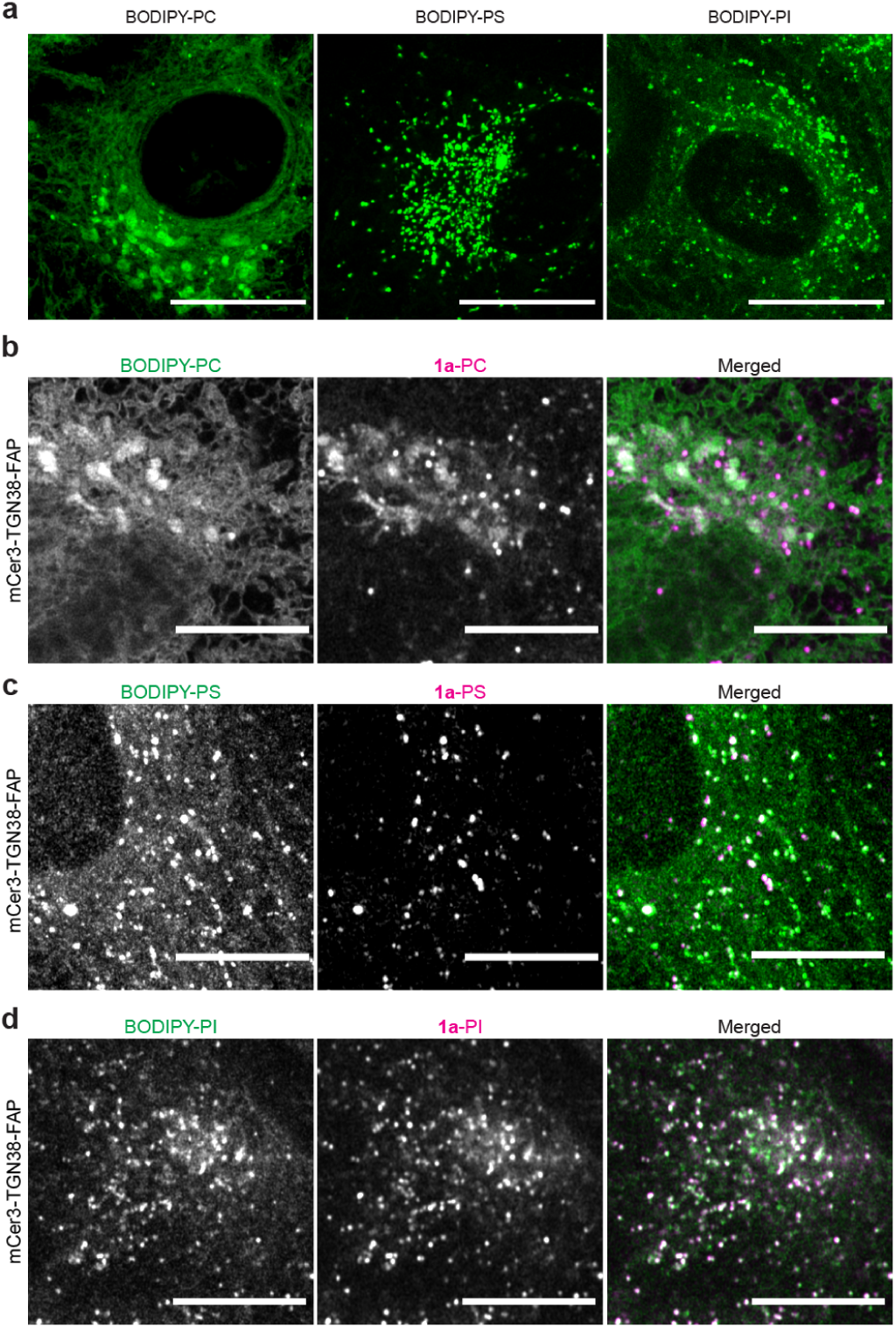
Multicolor imaging of global and targeted phospholipid pools. Cells were metabolically labeled with 100 μM N_3_-Cho, 250 μM C-L-Ser-N_3_, or 50 μM Ins-2-N_3_ for 24h prior to SPAAC conjugation. **a**, HeLa cells stained with 2 μM BODIPY-DBCO, which reacts with intracellular azido lipids. Images reflect maximal projection of Airyscan z-stacked images. Scale bars, 20 μm. **b**-**d**, Airyscan images of HeLa cells stably expressing mCer3-TGN38-FAP stained with equimolar **1a** and BODIPY-DBCO (1 μM each). Cells were incubated with 100 μM N_3_-Cho (**b**), 250 μM C-L-Ser-N_3_ (**c**), or 50 μM Ins-2-N_3_ for 24 h with doxycycline (1 mg/mL). 488 and 633 nm laser lines were used for imaging to avoid excitation of mCer3. Scale = 10 μm.

**Supplementary Fig. 11.**
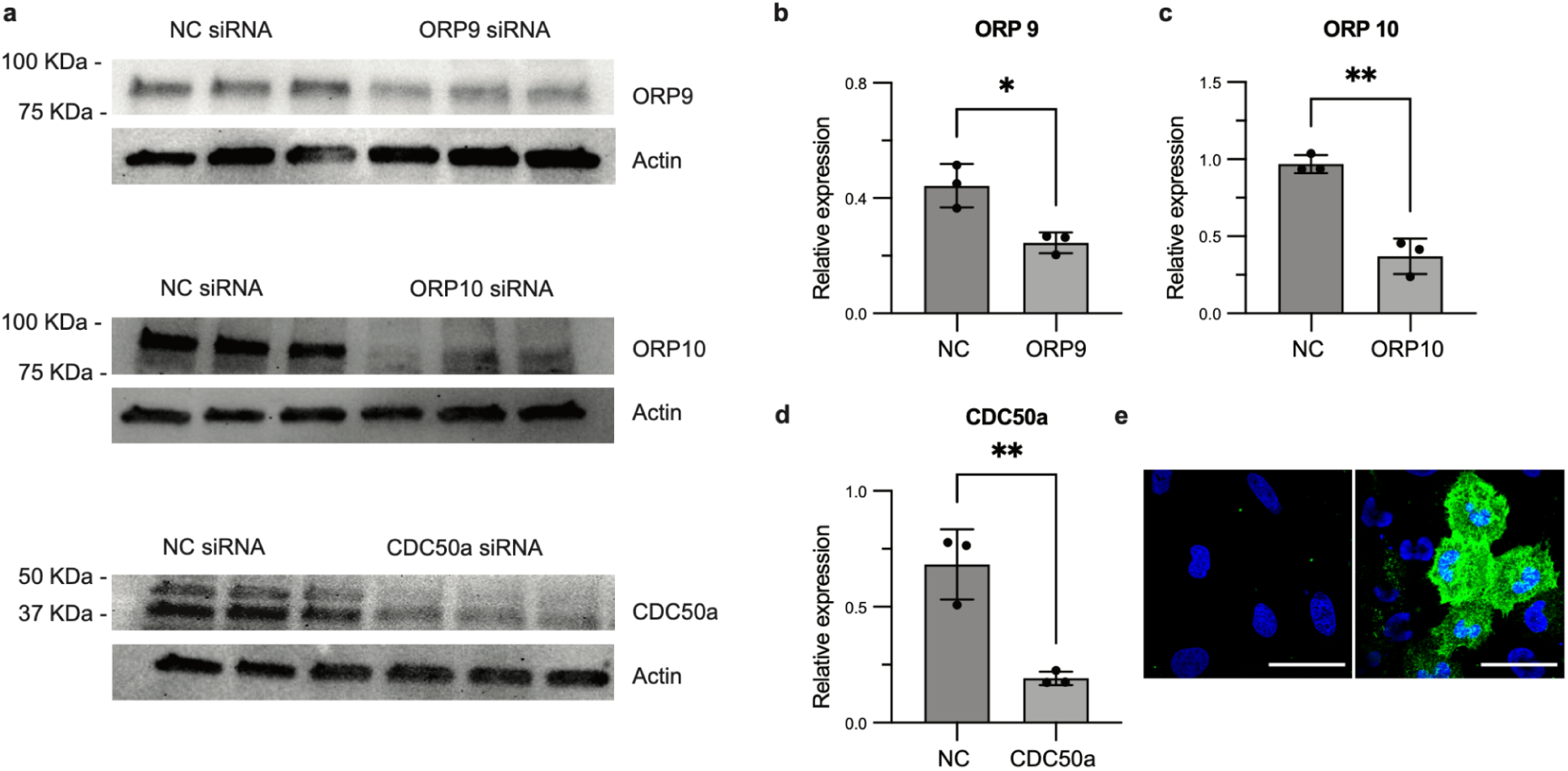
Silencing efficiency of ORP9, ORP10, and CDC50a siRNAs Cells were transfected with siRNA for 48 h prior to analysis. **a**, Western blot analysis of ORP9, ORP10, and CDC50A from silencing experiments. Antibodies for target genes were used at 1:1000 and actin was used as a loading control (1:5000). CDC50A and its actin control were blotted from two separate but identical SDS-PAGE gels run at the same time. **b**-**d**, Average band intensities normalized to actin (*n* = 3);Statistical significance was determined by Two-tailed Student’s T-test. **e**, Annexin V staining (green) of NC siRNA transfected cells (left) or CDC50A transfected cells (right). Nuclei were counterstained with Hoescht. Scale = 50 μm.

---

**Extended Data Fig. 1:** Mobility of TGN vesicles containing the soluble secretory cargo marker SP-mCherry Airyscan time-lapse of Golgi expressing SP-mCherry (magenta) and GFP-TGN38-FAP (green). Scale = 20 μm. https://drive.google.com/file/d/1TxF9G1RvI3twJ9pezZkijyKibNhc4T/view?usp=sharing

**Extended Data Fig. 2:** Mobility of TGN vesicles containing cytosolic PC and exoplasmic SM Airyscan time-lapse of Golgi from the cell shown in Fig. 4d expressing GFP-EqtSM (cyan) and mCer3-TGN38-FAP detecting **1a**-PC detection in the cytoplasmic leaflet (magenta). Images reflect maximal intensity values from z-stacked images spanning 1.5 μm. Scale = 10 μm. https://drive.google.com/file/d/1uWdM-LxW6N6hfmS_1oAvt9gelVmVm2tq/view?usp=sharing

**Extended Data Fig. 3:** Mobility of TGN vesicles containing cytosolic PC and protein cargoes Airyscan time-lapse of Golgi from the cell shown in Supplementary Fig. 10 expressing mCherry-GPI (yellow) and mCer3-TGN38-FAP detecting **1a**-PC detection in the cytoplasmic leaflet (magenta). Scale = 20 μm. https://drive.google.com/file/d/1S8RHj0p4P12DSThlM9wqXF7-yQTGs2qc/view?usp=sharing

## Supplementary Text: Synthesis of the Malachite Green Probes

Commercially available 4-nitrobenzaldehyde, *N*,*N*-dimethylaniline, niobium(V) chloride (NbCl_5_), 10% Pd/C, O-(7-azabenzotriazol-1-yl)-1,1,3,3-tetramethyluronium hexafluoro-phosphate (HATU), *N*,*N*-diisopropylethylamine (DIEA), tetrachloro-*p*-benzoquinone, and glacial acetic acid (AcOH) were obtained from Sigma-Aldrich. DBCO-C_6_-CO_2_H, azido-PEG_2_-CH_2_CO_2_H, and DBCO-PEG_2_-CO_2_H were obtained from BroadPharm. Deuterated chloroform (CDCl_3_) was obtained from Cambridge Isotope Laboratories. All reagents obtained from commercial suppliers were used without further purification unless otherwise noted. Analytical thin-layer chromatography was performed on E. Merck silica gel 60 F_254_ plates. Silica gel flash chromatography was performed using E. Merck silica gel (type 60SDS, 230-400 mesh). Solvent mixtures for chromatography are reported as v/v ratios. HPLC analysis was carried out on an Eclipse Plus C8 analytical column with *Phase A*/*Phase B* gradients [*Phase A*: H_2_O with 0.1% formic acid; *Phase B*: MeOH with 0.1% formic acid]. HPLC purification was carried out on Zorbax SB-C18 semipreparative column with *Phase A*/*Phase B* gradients [*Phase A*: H_2_O with 0.1% formic acid; *Phase B*: MeOH with 0.1% formic acid]. Proton nuclear magnetic resonance (^1^H NMR) spectra were recorded on a Jeol ECZL 400 MHz spectrometer, and were referenced relative to residual proton resonances in CDCl_3_ (at d 7.26 ppm) or CD_3_OD (at d 4.87 or 3.31 ppm). ^1^H NMR splitting patterns are assigned as singlet (s), doublet (d), triplet (t), quartet (q) or pentuplet (p). All first-order splitting patterns were designated on the basis of the appearance of the multiplet. Splitting patterns that could not be readily interpreted are designated as multiplet (m) or broad (br). Carbon nuclear magnetic resonance (^13^C NMR) spectra were recorded on a Jeol ECZL 400 MHz spectrometer, and were referenced relative to residual proton resonances in CDCl_3_ (at d 77.16 ppm) or CD_3_OD (at d 49.15 ppm). Electrospray Ionization-Time of Flight (ESI-TOF) spectra were obtained on an Agilent 6230 TOF-MS.

### 4,4’-((4-nitrophenyl)methylene)bis(*N*,*N*-dimethylaniline) [O_2_N-Malachite Green *(Leuco form)*]^77,78^

A mixture of 4-nitrobenzaldehyde (0.755 g, 5.0 mmol), *N*,*N*-dimethylaniline (1.90 mL, 15.0 mmol), and NbCl_5_ (0.135 g, 0.5 mmol) was heated at 130 °C for 2 h under Ar. Several color changes were observed, from yellow to orange, and finally purple. Afterwards, the reaction mixture was cooled to room temperature and diluted in EtOAc (20 mL) and H_2_O (20 mL). The aqueous layer was then extracted with EtOAc (3 × 10 mL), and the combined organic layers were dried over anhydrous MgSO_4_ and concentrated *in vacuo*. The crude was absorbed over SiO_2_, and then purified by flash column chromatography (10-20% EtOAc in hexanes), affording 1.486 g of O_2_N-Malachite Green *(Leuco form)* as an orange solid [79%, Rf = 0.42 (20% EtOAc in hexanes)]. ^1^H NMR (CDCl_3_, 400.13 MHz, d): 8.12 (d, *J* = 8.7 Hz, 2H, 2 × CH), 7.30 (d, *J* = 8.7 Hz, 2H, 2 × CH), 6.96 (d, *J* = 8.7 Hz, 4H, 4 × CH), 6.69 (d, *J* = 8.2 Hz, 4H, 4 × CH), 5.47 (s, 1H, 1 × CH), 2.94 (s, 12H, 4 × CH_3_). ^13^C NMR (CDCl_3_, 100.61 MHz, d): 153.3, 149.2, 146.3, 131.1, 130.2, 130.0, 123.6, 112.8, 55.0, 40.8. MS (ESI-TOF) [m/z (%)]: 376 ([MH]^+^, 34), 188 ([MH]^2+^, 100).

### 4,4’-((4-aminophenyl)methylene)bis(*N*,*N*-dimethylaniline) [H_2_N-Malachite Green *(Leuco form)*]

A suspension of 4,4’-((4-nitrophenyl)methylene)bis(*N*,*N*-dimethylaniline) [O_2_N-Malachite Green (*Leuco form*)), 150.0 mg, 0.40 mmol] and 10% Pd/C (42.6 mg) in absolute EtOH (8 mL) was stirred for 2 h under H_2_. Then, the mixture was filtered through a pad of celite, washing with EtOH (3 × 2 mL). Afterwards, the filtrate was concentrated *in vacuo*, affording 137.2 mg of H_2_N-Malachite Green (*Leuco form*) as a white solid [99%, Rf = 0.42 (20% EtOAc in hexanes)]. The product was used directly without any further purification. ^1^H NMR (CDCl_3_, 400.13 MHz, d): 7.00 (d, *J* = 8.7 Hz, 4H, 4 × CH), 6.92 (d, *J* = 8.3 Hz, 2H, 2 × CH), 6.69 (d, *J* = 8.7 Hz, 4H, 4 × CH), 6.61 (d, *J* = 8.3 Hz, 2H, 2 × CH), 5.29 (s, 1H, 1 × CH), 3.86 (br s, 2H, 1 × NH_2_), 2.92 (s, 12H, 4 × CH_3_). ^13^C NMR (CDCl_3_, 100.61 MHz, d): 148.8, 144.2, 135.8, 133.8, 130.2, 130.0, 115.2, 112.8, 54.2, 41.0. MS (ESI-TOF) [m/z (%)]: 346 ([MH]^+^, 11), 173 ([MH]^2+^, 100).

### DBCO-C_6_-Malachite Green (*Leuco form*)

A solution of DBCO-C_6_-CO_2_H (19.3 µL, 57.9 mmol) in 100 µL of CH_2_Cl_2_ was stirred at 0 °C for 10 min, and then HATU (24.2 mg, 63.7 mmol) and DIEA (40.4 µL, 231.6 µmol) were successively added. After 10 min stirring at 0 °C, a H_2_N-Malachite Green *(Leuco form)* (20.0 mg, 57.9 mmol) solution in 100 µL of CH_2_Cl_2_ containing DIEA (20.2 µL, 115.8 µmol) was added. After 1 h stirring at rt, the solvent was removed under reduced pressure to give a blue oil. The corresponding residue was dissolved in MeOH (400 µL), filtered using a 0.2 µm syringe-driven filter, and the crude solution was purified by HPLC, affording 29.2 mg of DBCO-C_6_-Malachite Green *(Leuco form)* as a blue oil [72%, t_R_ = 10.8 min (Zorbax SB-C18 semipreparative column, 50% *Phase A* in *Phase B*, 1 min, then 50-5% *Phase A* in *Phase B*, 5 min, and then 5% *Phase A* in *Phase B*, 10 min)]. ^1^H NMR (CDCl_3_, 400.13 MHz, d): 7.78 (s, 1H, 1 × NH), 7.72-7.64 (m, 1H, 1 × CH), 7.50-7.28 (m, 7H, 7 × CH), 7.25-7.18 (m, 4H, 4 × CH), 7.08-6.94 (m, 6H, 6 × CH), 6.93-6.72 (m, 2H, 2 × CH), 5.38 (s, 1H, 1 × CH), 5.17 (d, *J* = 13.8 Hz, 1H, 0.5 × CH_2_), 3.69 (d, *J* = 13.8 Hz, 1H, 0.5 × CH_2_), 2.96 (s, 12H, 4 × CH_3_), 2.29 (dt, *J_1_* = 16.0 Hz, *J_2_* = 6.0 Hz, 1H, 0.5 × CH_2_), 2.10 (dt, *J_1_* = 17.1 Hz, *J_2_* = 6.8 Hz, 1H, 0.5 × CH_2_), 2.02-1.81 (m, 2H, 1 × CH_2_), 1.62-1.40 (m, 3H, 1.5 × CH_2_), 1.39-1.21 (m, 1H, 0.5 × CH_2_). ^13^C NMR (CDCl_3_, 100.61 MHz, d): 173.8, 171.2, 151.7, 148.0, 132.5, 130.3, 129.8, 129.1, 128.6, 128.5, 128.4, 128.0, 127.3, 125.6, 123.1, 122.6, 119.9, 115.1, 108.0, 55.6, 54.7, 44.0, 36.8, 34.6, 25.1, 24.0. MS (ESI-TOF) [m/z (%)]: 661 ([MHl]^+^, 5), 331 ([MH]^2+^, 100).

### DBCO-C_6_-Malachite Green *(Chromatic form)* (**1a**)

To a blue solution of DBCO-C_6_-Malachite Green *(Leuco form)* (4.0 mg, 6.1 µmol) and tetrachloro-*p*-benzoquinone (1.8 mg, 7.3 µmol) in CHCl_3_ (226 µL), glacial acetic acid (4.5 µL, 0.45 µmol) was added. The solution was stirred at 35 °C for 2 h. Afterwards, the volatiles were removed under reduced pressure, obtaining a dark green residue. The crude was absorbed over SiO_2_, and then purified by flash column chromatography (0-7% MeOH in CH_2_Cl_2_), affording 3.8 mg of DBCO-C_6_-Malachite Green *(Chromatic form)* as a dark green solid [90%, Rf = 0.57 (10% MeOH in CH_2_Cl_2_)]. ^1^H NMR (CD_3_OD, 400.13 MHz, d): 7.91-7.76 (m, 4H, 4 × CH), 7.71-7.64 (m, 1H, 1 × NH), 7.57-7.47 (m, 2H, 2 × CH), 7.44 (d, *J* = 9.0 Hz, 4H, 4 × CH), 7.37 (d, *J* = 8.7 Hz, 2H, 2 × CH), 7.33-7.19 (m, 4H, 4 × CH), 7.06 (d, *J* = 9.4 Hz, 4H, 4 × CH), 4.93-4.91 (m, 2H, 1 × CH_2_), 3.33 (s, 12H, 4 × CH_3_), 2.52-2.45 (m, 2H, 1 × CH_2_), 2.44-2.34 (m, 2H, 1 × CH_2_), 1.83-1.65 (m, 4H, 2 × CH_2_). ^13^C NMR (CD_3_OD, 100.61 MHz, d): 178.9, 174.7, 158.4, 145.6, 141.9, 137.7, 135.8, 135.6, 134.2, 133.5, 130.8, 130.4, 130.0, 129.7, 129.2, 128.9, 128.8, 128.3, 126.5, 125.6, 124.3, 123.1, 120.7, 120.2, 114.8, 114.5, 114.2, 40.9, 37.7, 34.4, 33.1, 30.8, 26.0, 25.5. MS (ESI-TOF) [m/z (%)]: 660 ([MH-Cl]^+^, 51), 659 ([M-Cl]^+^, 100), 330 ([MH-Cl]^2+^, 53).

### DBCO-PEG_2_-Malachite Green *(Leuco form)*

A solution of DBCO-PEG_2_-CO_2_H (13.4 mg, 28.9 mmol) in 200 µL of CH_2_Cl_2_/DMF (1:1) was stirred at 0 °C for 10 min, and then HATU (12.1 mg, 31.8 mmol) and DIEA (20.2 µL, 115.8 µmol) were successively added. After 10 min stirring at 0 °C, a H_2_N-Malachite Green *(Leuco form)* (10.0 mg, 28.9 mmol) solution in 200 µL of CH_2_Cl_2_/DMF (1:1) containing DIEA (10.1 µL, 57.9 µmol) was added. After 1 h stirring at rt, the solvent was removed under reduced pressure to give a blue oil. The corresponding residue was dissolved in MeOH (200 µL), filtered using a 0.2 µm syringe-driven filter, and the crude solution was purified by HPLC, affording 13.2 mg of DBCO-PEG_2_-Malachite Green *(Leuco form)* as a blue oil [55%, t_R_ = 50.9 min (Zorbax SB-C18 semipreparative column, 95% *Phase A* in *Phase B*, 1 min, then 95-50% *Phase A* in *Phase B*, 30 min, and then 50-0% *Phase A* in *Phase B*, 22 min)]. ^1^H NMR (CDCl_3_, 400.13 MHz, d): 8.76 (s, 1H, 1 × NH), 7.60-7.54 (m, 1H, 1 × CH), 7.53-7.46 (m, 1H, 1 × CH), 7.46-7.34 (m, 6H, 6 × CH), 7.25-7.17 (m, 4H, 4 × CH), 7.09-6.96 (m, 6H, 6 × CH), 6.95-6.72 (m, 2H, 2 × CH), 6.28 (t, *J* = 5.6 Hz, 1H, 1 × NH), 5.36 (s, 1H, 1 × CH), 5.09 (d, *J* = 13.9 Hz, 1H, 0.5 × CH_2_), 3.87-3.71 (m, 2H, 1 × CH_2_), 3.69-3.56 (m, 5H, 2.5 × CH_2_), 3.45 (t, *J* = 5.4 Hz, 2H, 1 × CH_2_), 3.34-3.18 (m, 2H, 1 × CH_2_), 2.95 (s, 12H, 4 × CH_3_), 2.87-2.75 (m, 1H, 0.5 × CH_2_), 2.53-2.41 (m, 2H, 1 × CH_2_), 2.39-2.26 (m, 1H, 0.5 × CH_2_), 2.17-2.02 (m, 1H, 0.5 × CH_2_), 1.98-1.83 (m, 1H, 0.5 × CH_2_). ^13^C NMR (CDCl_3_, 100.61 MHz, d): 172.7, 172.1, 169.9, 151.3, 148.1, 132.3, 130.3, 129.8, 129.5, 128.9, 128.4, 128.2, 127.9, 127.2, 125.6, 123.2, 122.6, 119.9, 114.8, 108.0, 70.7, 69.9, 69.8, 67.4, 55.8, 54.8, 39.1, 38.0, 31.5, 30.3. MS (ESI-TOF) [m/z (%)]: 793 ([MH]^+^, 4), 397 ([MH]^2+^, 100).

### DBCO-PEG_2_-Malachite Green *(Chromatic form)* (**1b**)

To a blue solution of DBCO-PEG_2_-Malachite Green *(Leuco form)* (5.0 mg, 6.3 µmol) and tetrachloro-*p*-benzoquinone (1.9 mg, 7.6 µmol) in CHCl_3_ (235 µL), glacial acetic acid (4.7 µL, 0.47 µmol) was added. The solution was stirred at 35 °C for 2 h. Afterwards, the volatiles were removed under reduced pressure, obtaining a dark green residue. The crude was absorbed over SiO_2_, and then purified by flash column chromatography (0-10% MeOH in CH_2_Cl_2_), affording 4.8 mg of N_3_-PEG_2_-Malachite Green *(Chromatic form)* as a dark green solid [92%, Rf = 0.44 (10% MeOH in CH_2_Cl_2_)]. ^1^H NMR (CDCl_3_, 400.13 MHz, d): 11.33 (s, 1H, 1 × NH), 8.20 (d, *J* = 8.4 Hz, 2H, 2 × CH), 7.85-7.69 (m, 1H, 1 × NH), 7.63 (d, *J* = 7.3 Hz, 1H, 1 × CH), 7.58 (d, *J* = 7.6 Hz, 1H, 1 × CH), 7.42-7.27 (m, 9H, 9 × CH), 7.25-7.17 (m, 3H, 3 × CH), 6.86 (d, *J* = 8.5 Hz, 4H, 4 × CH), 5.13 (d, *J* = 13.9 Hz, 1H, 0.5 × CH_2_), 3.91 (t, *J* = 5.9 Hz, 2H, 1 × CH_2_), 3.76-3.57 (m, 5H, 2.5 × CH_2_), 3.56-3.45 (m, 2H, 1 × CH_2_), 3.31 (s, 12H, 4 × CH_3_), 2.94 (t, *J* = 5.9 Hz, 2H, 1 × CH_2_), 2.87-2.75 (m, 1H, 0.5 × CH_2_), 2.68-2.53 (m, 1H, 0.5 × CH_2_), 2.46-2.26 (m, 4H, 2 × CH_2_). ^13^C NMR (CDCl_3_, 100.61 MHz, d): 175.7, 174.4, 172.4, 156.5, 146.3, 141.1, 137.4, 134.2, 133.4, 133.3, 132.3, 129.3, 127.7, 127.6, 125.8, 124.9, 124.4, 122-1, 121.9, 120.0, 119.9, 117.3, 113.6, 133.5, 98.1, 98.0, 70.4, 70.3, 68.9, 67.3, 47.0, 41.2, 40.4, 38.0, 32.1, 29.9, 29.5, 22.9*, 14.3*. MS (ESI-TOF) [m/z (%)]: 791 ([MH-Cl]^+^, 3), 396 ([MH-Cl]^2+^, 100). *Artifact.

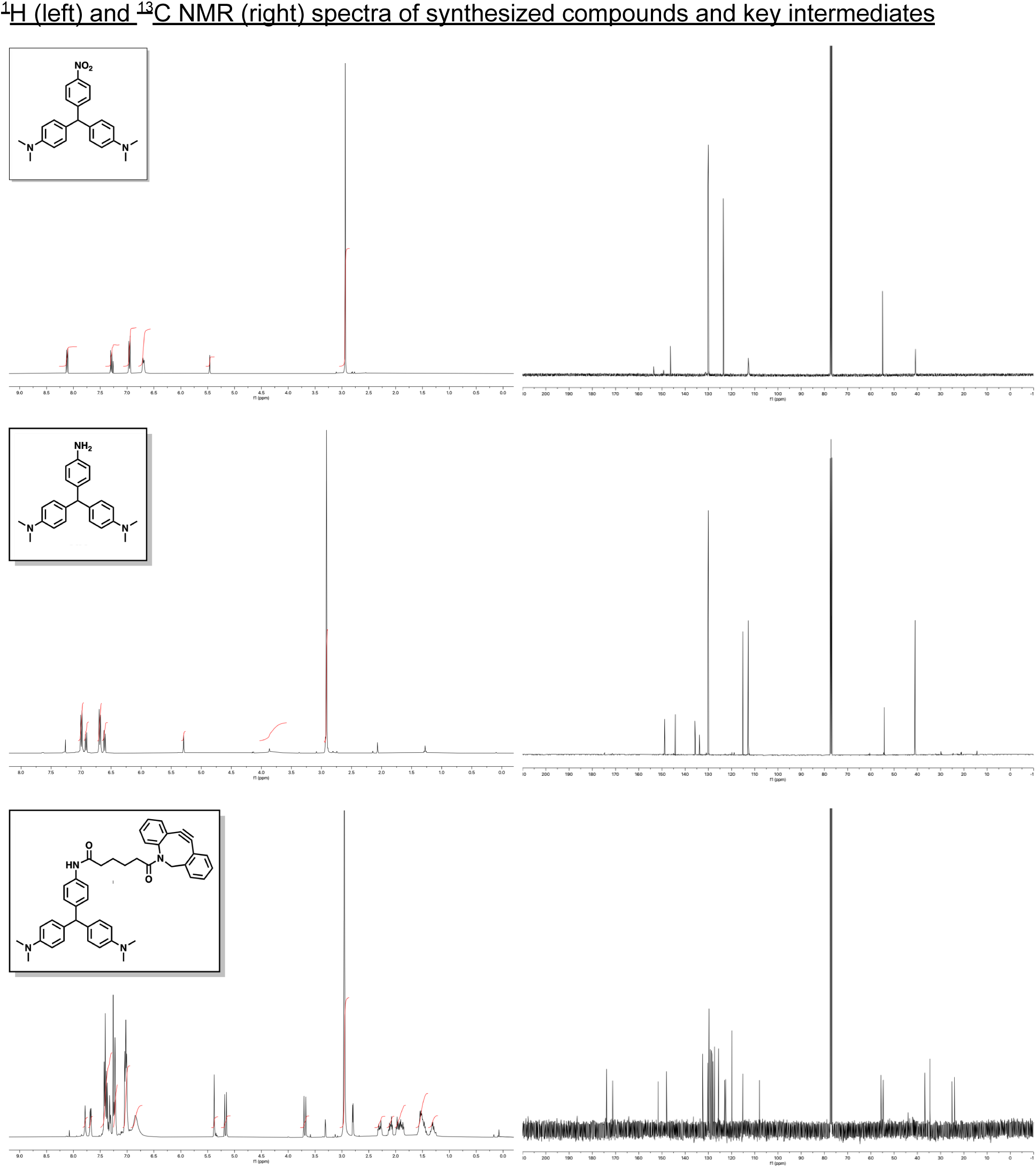

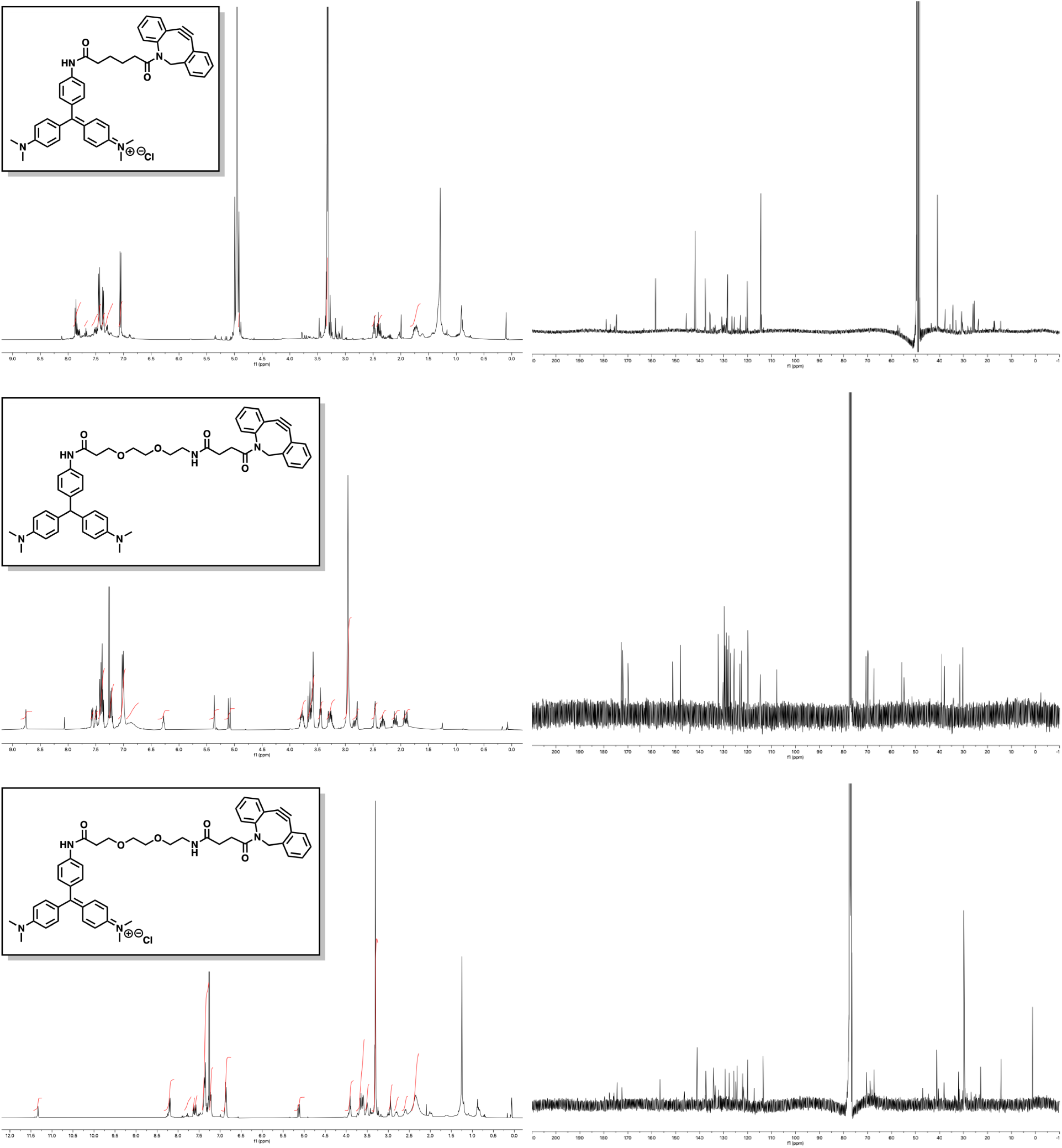

## References

1. van Meer, G., Voelker, D. R. & Feigenson, G. W. Membrane lipids: where they are and how they behave. Nat. Rev. Mol. Cell Biol. 9, 112–124 (2008).

2. Levental, I. & Lyman, E. Regulation of membrane protein structure and function by their lipid nano-environment. Nat. Rev. Mol. Cell Biol. 24, 107–122 (2023).

3. Lorent, J. H. et al. Plasma membranes are asymmetric in lipid unsaturation, packing and protein shape. Nat. Chem. Biol. 16, 644–652 (2020).

4. Khaddaj, R. & Kukulski, W. Piecing together the structural organisation of lipid exchange at membrane contact sites. Curr. Opin. Cell Biol. 83, 102212 (2023).

5. Voeltz, G. K., Sawyer, E. M., Hajnóczky, G. & Prinz, W. A. Making the connection: How membrane contact sites have changed our view of organelle biology. Cell 187, 257–270 (2024).

6. Wong, L. H., Gatta, A. T. & Levine, T. P. Lipid transfer proteins: the lipid commute via shuttles, bridges and tubes. Nat. Rev. Mol. Cell Biol. 20, 85–101 (2019).

7. Lipp, N.-F., Ikhlef, S., Milanini, J. & Drin, G. Lipid Exchangers: Cellular Functions and Mechanistic Links With Phosphoinositide Metabolism. Front Cell Dev Biol 8, 663 (2020).

8. Rothman, J. E. & Lenard, J. Membrane asymmetry. Science 195, 743–753 (1977).

9. Verkleij, A. J. et al. The asymmetric distribution of phospholipids in the human red cell membrane. A combined study using phospholipases and freeze-etch electron microscopy. Biochim. Biophys. Acta 323, 178–193 (1973).

10. Jao, C. Y., Roth, M., Welti, R. & Salic, A. Metabolic labeling and direct imaging of choline phospholipids in vivo. Proc. Natl. Acad. Sci. U. S. A. 106, 15332–15337 (2009).

11. Ancajas, C. F. et al. Cellular labeling of phosphatidylserine using clickable Serine probes. ACS Chem. Biol. 18, 377–384 (2023).

12. Ricks, T. J. et al. Labeling of Phosphatidylinositol Lipid Products in Cells through Metabolic Engineering by Using a Clickable myo-Inositol Probe. Chembiochem 20, 172–180 (2019).

13. Bumpus, T. W. & Baskin, J. M. A Chemoenzymatic Strategy for Imaging Cellular Phosphatidic Acid Synthesis. Angew. Chem. Int. Ed Engl. 55, 13155–13158 (2016).

14. Bussink, A. P. et al. N-azidoacetylmannosamine-mediated chemical tagging of gangliosides. J. Lipid Res. 48, 1417–1421 (2007).

15. Tamura, T. et al. Organelle membrane-specific chemical labeling and dynamic imaging in living cells. Nat. Chem. Biol. 16, 1361–1367 (2020).

16. Maekawa, M., Yang, Y. & Fairn, G. D. Perfringolysin O Theta Toxin as a Tool to Monitor the Distribution and Inhomogeneity of Cholesterol in Cellular Membranes. Toxins 8, (2016).

17. Deng, Y., Rivera-Molina, F. E., Toomre, D. K. & Burd, C. G. Sphingomyelin is sorted at the trans Golgi network into a distinct class of secretory vesicle. Proc. Natl. Acad. Sci. U. S. A. 113, 6677–6682 (2016).

18. Gallo, E. Fluorogen-Activating Proteins: Next-Generation Fluorescence Probes for Biological Research. Bioconjug. Chem. 31, 16–27 (2020).

19. Kozma, E. & Kele, P. Fluorogenic probes for super-resolution microscopy. Org. Biomol. Chem. 17, 215–233 (2019).

20. Bottone, S. et al. A fluorogenic chemically induced dimerization technology for controlling, imaging and sensing protein proximity. Nat. Methods 20, 1553–1562 (2023).

21. Yan, Q. et al. Localization microscopy using noncovalent fluorogen activation by genetically encoded fluorogen-activating proteins. Chemphyschem 15, 687–695 (2014).

22. Horibata, Y. & Sugimoto, H. Differential contributions of choline phosphotransferases CPT1 and CEPT1 to the biosynthesis of choline phospholipids. J. Lipid Res. 62, 100100 (2021).

23. Liu, T. et al. Multi-color live-cell STED nanoscopy of mitochondria with a gentle inner membrane stain. Proc. Natl. Acad. Sci. U. S. A. 119, e2215799119 (2022).

24. Stephan, T., Roesch, A., Riedel, D. & Jakobs, S. Live-cell STED nanoscopy of mitochondrial cristae. Sci. Rep. 9, 12419 (2019).

25. John Peter, A. T., Petrungaro, C., Peter, M. & Kornmann, B. METALIC reveals interorganelle lipid flux in live cells by enzymatic mass tagging. Nat. Cell Biol. 24, 996–1004 (2022).

26. Kumar, N. et al. VPS13A and VPS13C are lipid transport proteins differentially localized at ER contact sites. J. Cell Biol. 217, 3625–3639 (2018).

27. Horibata, Y. et al. StarD7 Protein Deficiency Adversely Affects the Phosphatidylcholine Composition, Respiratory Activity, and Cristae Structure of Mitochondria. J. Biol. Chem. 291, 24880–24891 (2016).

28. Horibata, Y. & Sugimoto, H. StarD7 mediates the intracellular trafficking of phosphatidylcholine to mitochondria. J. Biol. Chem. 285, 7358–7365 (2010).

29. Horibata, Y. et al. Identification of the N-terminal transmembrane domain of StarD7 and its importance for mitochondrial outer membrane localization and phosphatidylcholine transfer. Sci. Rep. 7, 8793 (2017).

30. Bos, K., Wraight, C. & Stanley, K. K. TGN38 is maintained in the trans-Golgi network by a tyrosine-containing motif in the cytoplasmic domain. EMBO J. 12, 2219–2228 (1993).

31. Luzio, J. P. et al. Identification, sequencing and expression of an integral membrane protein of the trans-Golgi network (TGN38). Biochem. J 270, 97–102 (1990).

32. Lieu, Z. Z. & Gleeson, P. A. Identification of different itineraries and retromer components for endosome-to-Golgi transport of TGN38 and Shiga toxin. Eur. J. Cell Biol. 89, 379–393 (2010).

33. Iyoshi, S. et al. Asymmetrical distribution of choline phospholipids revealed by click chemistry and freeze-fracture electron microscopy. ACS Chem. Biol. 9, 2217–2222 (2014).

34. Cabukusta, B. et al. A lipid transfer protein knockout library reveals ORP9-ORP11 dimer mediating PS/PI(4)P exchange at the ER-trans Golgi contact site to promote sphingomyelin synthesis. bioRxiv 2023.06.02.543249 (2023) doi:10.1101/2023.06.02.543249.

35. Phillips, S. E., Ile, K. E., Boukhelifa, M., Huijbregts, R. P. H. & Bankaitis, V. A. Specific and nonspecific membrane-binding determinants cooperate in targeting phosphatidylinositol transfer protein beta-isoform to the mammalian trans-Golgi network. Mol. Biol. Cell 17, 2498–2512 (2006).

36. Fairn, G. D. et al. High-resolution mapping reveals topologically distinct cellular pools of phosphatidylserine. J. Cell Biol. 194, 257–275 (2011).

37. Saxon, E. & Bertozzi, C. R. Cell surface engineering by a modified Staudinger reaction. Science 287, 2007–2010 (2000).

38. Fleischer, B. Mechanism of glycosylation in the Golgi apparatus. J. Histochem. Cytochem. 31, 1033–1040 (1983).

39. Klemm, R. W. et al. Segregation of sphingolipids and sterols during formation of secretory vesicles at the trans-Golgi network. J. Cell Biol. 185, 601–612 (2009).

40. Doktorova, M. et al. Cell Membranes Sustain Phospholipid Imbalance Via Cholesterol Asymmetry. bioRxiv 2023.07.30.551157 (2023) doi:10.1101/2023.07.30.551157.

41. Takar, M., Huang, Y. & Graham, T. R. The PQ-loop protein Any1 segregates Drs2 and Neo1 functions required for viability and plasma membrane phospholipid asymmetry. J. Lipid Res. 60, 1032–1042 (2019).

42. Hankins, H. M., Sere, Y. Y., Diab, N. S., Menon, A. K. & Graham, T. R. Phosphatidylserine translocation at the yeast trans-Golgi network regulates protein sorting into exocytic vesicles. Mol. Biol. Cell 26, 4674–4685 (2015).

43. Takar, M., Wu, Y. & Graham, T. R. The essential Neo1 protein from budding yeast plays a role in establishing aminophospholipid asymmetry of the plasma membrane. J. Biol. Chem. 291, 15727–15739 (2016).

44. Chen, S. et al. Roles for the Drs2p-Cdc50p complex in protein transport and phosphatidylserine asymmetry of the yeast plasma membrane. Traffic 7, 1503–1517 (2006).

45. Alder-Baerens, N., Lisman, Q., Luong, L., Pomorski, T. & Holthuis, J. C. M. Loss of P4 ATPases Drs2p and Dnf3p disrupts aminophospholipid transport and asymmetry in yeast post-Golgi secretory vesicles. Mol. Biol. Cell 17, 1632–1642 (2006).

46. He, R. et al. ORP9 and ORP10 form a heterocomplex to transfer phosphatidylinositol 4-phosphate at ER-TGN contact sites. Cell. Mol. Life Sci. 80, 77 (2023).

47. Kawasaki, A. et al. PI4P/PS countertransport by ORP10 at ER-endosome membrane contact sites regulates endosome fission. J. Cell Biol. 221, (2022).

48. Cabukusta, B. et al. A lipid transfer protein knockout library reveals ORP9-ORP11 dimer mediating PS/PI(4)P exchange at the ER-trans Golgi contact site to promote sphingomyelin synthesis. eLife (2023) doi:10.7554/elife.91345.1.

49. Tanaka, Y. et al. The phospholipid flippase ATP9A is required for the recycling pathway from the endosomes to the plasma membrane. Mol. Biol. Cell 27, 3883–3893 (2016).

50. Takatsu, H. et al. Phospholipid Flippase Activities and Substrate Specificities of Human Type IV P-type ATPases Localized to the Plasma Membrane *. J. Biol. Chem. 289, 33543–33556 (2014).

51. Best, J. T., Xu, P. & Graham, T. R. Phospholipid flippases in membrane remodeling and transport carrier biogenesis. Curr. Opin. Cell Biol. 59, 8–15 (2019).

52. Segawa, K., Kurata, S. & Nagata, S. The CDC50A extracellular domain is required for forming a functional complex with and chaperoning phospholipid flippases to the plasma membrane. J. Biol. Chem. 293, 2172–2182 (2018).

53. Coleman, J. A. & Molday, R. S. Critical role of the beta-subunit CDC50A in the stable expression, assembly, subcellular localization, and lipid transport activity of the P4-ATPase ATP8A2. J. Biol. Chem. 286, 17205–17216 (2011).

54. Boyce, M. et al. Metabolic cross-talk allows labeling of O-linked β-*N*-acetylglucosamine-modified proteins via the *N*-acetylgalactosamine salvage pathway. Proceedings of the National Academy of Sciences 108, 3141–3146 (2011).

55. Hang, H. C., Yu, C., Kato, D. L. & Bertozzi, C. R. A metabolic labeling approach toward proteomic analysis of mucin-type O-linked glycosylation. Proc. Natl. Acad. Sci. U. S. A. 100, 14846–14851 (2003).

56. Ma, Z. & Vosseller, K. Cancer Metabolism and Elevated O-GlcNAc in Oncogenic Signaling*. J. Biol. Chem. 289, 34457–34465 (2014).

57. Tan, E. P. et al. Sustained O-GlcNAcylation reprograms mitochondrial function to regulate energy metabolism. J. Biol. Chem. 292, 14940–14962 (2017).

58. Zhu, Q. et al. *O*-GlcNAcylation promotes tumor immune evasion by inhibiting PD-L1 lysosomal degradation. Proceedings of the National Academy of Sciences 120, e2216796120 (2023).

59. Pekkurnaz, G., Trinidad, J. C., Wang, X., Kong, D. & Schwarz, T. L. Glucose regulates mitochondrial motility via Milton modification by O-GlcNAc transferase. Cell 158, 54–68 (2014).

60. Dupas, T. et al. An overview of tools to decipher O-GlcNAcylation from historical approaches to new insights. Int. J. Biochem. Cell Biol. 151, 106289 (2022).

61. Onodera, Y., Nam, J.-M. & Bissell, M. J. Increased sugar uptake promotes oncogenesis via EPAC/RAP1 and O-GlcNAc pathways. J. Clin. Invest. 124, 367–384 (2014).

62. Jóźwiak, P. et al. Mitochondrial O-GlcNAc Transferase Interacts with and Modifies Many Proteins and Its Up-Regulation Affects Mitochondrial Function and Cellular Energy Homeostasis. Cancers 13, (2021).

63. Ma, J. et al. O-GlcNAcomic Profiling Identifies Widespread O-Linked β-N-Acetylglucosamine Modification (O-GlcNAcylation) in Oxidative Phosphorylation System Regulating Cardiac Mitochondrial Function. J. Biol. Chem. 290, 29141–29153 (2015).

64. Guzmán, L. E. et al. Bioorthogonal Metabolic Labeling of the Virulence Factor Phenolic Glycolipid in Mycobacteria. ACS Chem. Biol. (2024) doi:10.1021/acschembio.3c00724.

65. Fantoni, N. Z., El-Sagheer, A. H. & Brown, T. A Hitchhiker’s Guide to Click-Chemistry with Nucleic Acids. Chem. Rev. 121, 7122–7154 (2021).

66. Liang, D. et al. A real-time, click chemistry imaging approach reveals stimulus-specific subcellular locations of phospholipase D activity. Proc. Natl. Acad. Sci. U. S. A. 116, 15453–15462 (2019).

67. Nguyen, S. S. & Prescher, J. A. Developing bioorthogonal probes to span a spectrum of reactivities. Nat Rev Chem 4, 476–489 (2020).

68. Sakuragi, T. & Nagata, S. Regulation of phospholipid distribution in the lipid bilayer by flippases and scramblases. Nat. Rev. Mol. Cell Biol. 24, 576–596 (2023).

69. Surma, M. A., Klose, C. & Simons, K. Lipid-dependent protein sorting at the trans-Golgi network. Biochim. Biophys. Acta 1821, 1059–1067 (2012).

70. Sharpe, H. J., Stevens, T. J. & Munro, S. A comprehensive comparison of transmembrane domains reveals organelle-specific properties. Cell 142, 158–169 (2010).

71. Carpenter, M. A. et al. Protein Proximity Observed Using Fluorogen Activating Protein and Dye Activated by Proximal Anchoring (FAP-DAPA) System. ACS Chem. Biol. 15, 2433–2443 (2020).

72. Telmer, C. A. et al. Rapid, specific, no-wash, far-red fluorogen activation in subcellular compartments by targeted fluorogen activating proteins. ACS Chem. Biol. 10, 1239–1246 (2015).

73. Kowarz, E., Löscher, D. & Marschalek, R. Optimized Sleeping Beauty transposons rapidly generate stable transgenic cell lines. Biotechnol. J. 10, 647–653 (2015).

74. Jao, C. Y., Roth, M., Welti, R. & Salic, A. Biosynthetic labeling and two-color imaging of phospholipids in cells. Chembiochem 16, 472–476 (2015).

75. Sparkes, B. L., Slone, E. E. A., Roth, M., Welti, R. & Fleming, S. D. Intestinal lipid alterations occur prior to antibody-induced prostaglandin E2 production in a mouse model of ischemia/reperfusion. Biochim. Biophys. Acta 1801, 517–525 (2010).

76. Zhou, Z. et al. LipidomeDB data calculation environment: online processing of direct-infusion mass spectral data for lipid profiles. Lipids 46, 879–884 (2011).

77. Montagut, A. M., Gálvez, E., Shafir, A., Sebastián, R. M. & Vallribera, A. Triarylmethane Dyes for Artificial Repellent Cotton Fibers. Chemistry 23, 3810–3814 (2017).

78. Kim, J., Park, E. Y., Moon, S.-Y., Kim, J.-M. & Park, K. Patterned color images with a triphenylmethane-derived acrylate polymer. Macromol. Res. 16, 81–83 (2008).

